# Decomposing leaf mass into metabolic and structural components explains divergent patterns of trait variation within and among plant species

**DOI:** 10.1101/116855

**Authors:** Masatoshi Katabuchi, Kaoru Kitajima, S. Joseph Wright, Sunshine A. Van Bael, Jeanne L. D. Osnas, Jeremy W. Lichstein

**Author notes:** Corresponding Author:; Masatoshi Katabuchi: https://orcid.org/0000-0001-9900-9029 Kaoru Kitajima: https://orcid.org/0000-0001-6822-8536 S. Joseph Wright: https://orcid.org/0000-0003-4260-5676 Sunshine A. Van Bael: https://orcid.org/0000-0001-7317-3533 Jeremy W. Lichstein: https://orcid.org/0000-0001-5553-6142.

## Abstract

1. Across the global flora, interspecific variation in photosynthetic and metabolic rates depends more strongly on leaf area than leaf mass. In contrast, intraspecific variation in these rates is strongly mass-dependent. These contrasting patterns suggest that the causes of variation in leaf mass per area (LMA) may be fundamentally different within vs. among species.
2. We developed a statistical modeling framework to decompose LMA into two conceptual components – metabolic LMAm (which determines photosynthetic capacity and dark respiration) and structural LMAs (which determines leaf toughness and potential leaf lifespan) - using leaf trait data from tropical forests in Panama and a global leaf-trait database.
3. Decomposing LMA into LMAm and LMAs improves predictions of leaf trait variation (photosynthesis, respiration, and lifespan). We show that strong area-dependence of metabolic traits across species can result from multiple factors, including high LMAs variance and/or a slow increase in photosynthetic capacity with increasing LMAm. In contrast, strong mass-dependence of metabolic traits within species results from LMAm increasing from sunny to shady conditions. LMAm and LMAs were nearly independent of each other in both global and Panama datasets.
4. *Synthesis*: Our results suggest that leaf functional variation is multi-dimensional and that biogeochemical models should treat metabolic and structural leaf components separately.

## Introduction

Leaf functional traits play an important role in ecological and physiological tradeoffs (Wright et al. 2004, Reich 2014, Onoda et al. 2017) and in carbon and nutrient cycling (Tcherkez et al. 2017, e.g., Huntingford et al. 2017). Thus, understanding the causes and consequences of leaf trait variation is an important goal in ecology, plant physiology, and biogeochemistry (Bonan et al. 2002, Poorter et al. 2009). Different leaf assemblages exhibit markedly different patterns of trait variation. For example, across global species, whole-leaf values of traits related to photosynthesis and metabolism (e.g., the photosynthetic capacity, respiration rate, or nutrient content of entire leaves) tend to increase more strongly with leaf area than leaf mass (Osnas et al. 2013), whereas the opposite pattern (strong mass-dependence of these same traits) is observed within species across light gradients (Osada et al. 2001, Osnas et al. 2018). Functional groups (e.g., deciduous vs. evergreen angiosperms) also differ from each other in terms of how photosynthetic and metabolic traits variation depend on leaf mass and area (Osnas et al. 2018). These divergent patterns suggest the presence of multiple drivers of trait variation.

Strong interspecific correlations among leaf mass per area (LMA), leaf lifespan (LL), and mass-normalized leaf traits related to photosynthesis and metabolism (hereafter, ‘metabolic traits’) have been interpreted as evidence for a single dominant axis of leaf functional variation (Wright et al. 2004). However, the interpretation of these strong correlations, which can result from mass-normalization of area-dependent traits (Lloyd et al. 2013, Osnas et al. 2013), is controversial. On the one hand, mass-normalization has been justified based on economic principles alone (Westoby et al. 2013), because leaf mass provides a simple index of investment that underlies the leaf economics spectrum (LES), ranging from cheap, low-LMA leaves with a fast rate of return per-unit investment to expensive, high-LMA leaves with a slow rate of return (Wright et al. 2004). On the other hand, because the lifetime return on investment depends on LL (Westoby et al. 2000, Falster et al. 2012), traits should be normalized by their annualized construction costs (or mass/LL) should be used as a normalizer in leaf economics analyses (Osnas et al. 2013). In global analyses, normalizing leaf traits by mass/LL yields similarly weak correlations as area-normalization, because leaf area is roughly proportional to mass/LL across global species (Osnas et al. 2013). Area-dependence of metabolic traits and the inconsistent correlation strengths obtained from different normalizers (Lloyd et al. 2013, Osnas et al. 2013) do not invalidate mass-normalization or the LES (Westoby et al. 2013). These considerations do, however, lead us to question the evidence for a single dominant axis of leaf functional diversity, which has important implications for how trait variation is represented in ecosystem models (Bonan et al. 2002, Sakschewski et al. 2016).

To understand why the same data can be interpreted as either supporting or opposing the existence of a single dominant axis of leaf trait variation, consider the conceptual model proposed by Osnas et al. (2018), in which LMA is comprised of two additive components: metabolic LMA (LMAm) – the mass per area of chloroplasts and other metabolically active leaf components that contribute directly to photosynthesis and respiration – and structural LMA (LMAs) – the mass per area of structural leaf components that contribute to toughness and durability (Kitajima et al. 2012, 2016, Onoda et al. 2017). Suppose these two LMA components are independent axes of functional variation, so that LMAm and LMAs are uncorrelated across species (Fig. 1a). These two independent axes can be translated into either a two-dimensional trait space (if metabolic traits are area-normalized; Fig. 1b) or a one-dimensional trait space (if metabolic traits are mass-normalized; Fig. 1c). While both Fig. 1b and Fig. 1c are both ‘correct’ representations of the same data, they lead to different perceptions about the dimensionality of leaf functional variation. If LMAm, LMAs, and/or other axes are largely independent and have distinct functional consequences, then it would not be possible to accurately represent functional variation with a single axis.

**Fig. 1:**
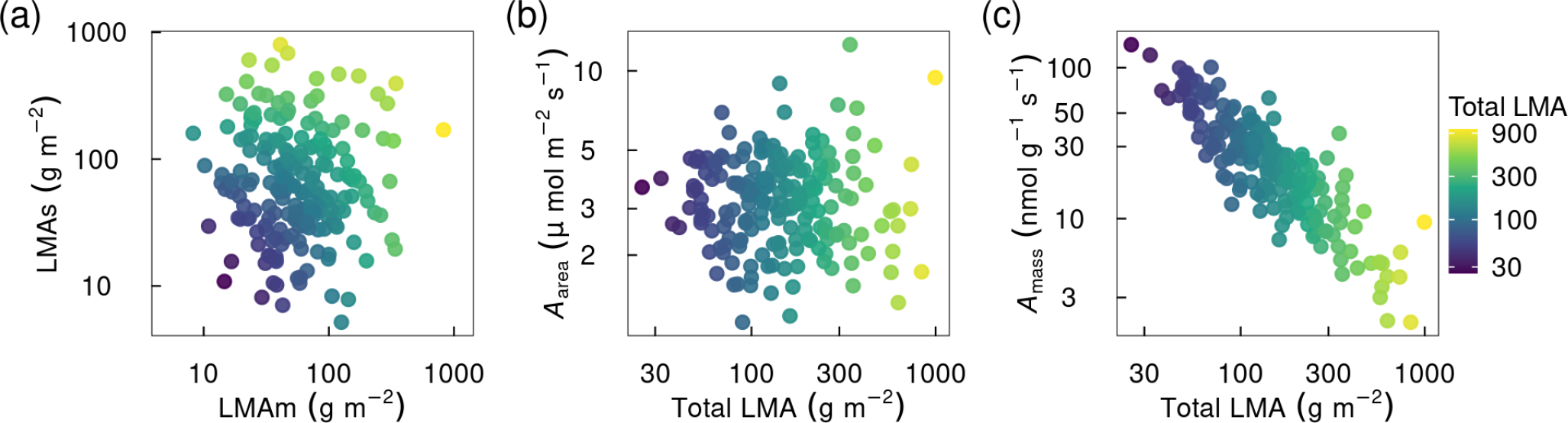
Simulated two-dimensional functional space that can result in either a two- or one-dimensional trait space, depending on how metabolic traits are normalized. (a) Hypothetical independent variation in two leaf mass per area (LMA) components: metabolic leaf mass per area (LMAm, which largely determines per-area values of photosynthesis, respiration, and nutrient concentrations) and structural leaf mass per area (LMAs, which determines leaf toughness but has little effect on metabolic traits). Panels b and c represent normalization by leaf area and leaf mass, respectively. (b) Two-dimensional relationship between photosynthetic capacity (*A*_max_) per-unit leaf area and total LMA (equal to LMAm + LMAs). (c) One-dimensional relationship between *A*_max_ per-unit leaf mass and total LMA. Methods used to simulate the data are explained in Appendix S3.

The hypothetical example in Fig. 1 shows how mass-normalization can, in principle, make a two-dimensional trait space appear one-dimensional, but the dimensionality of functional variation in real leaf assemblages remains an open question. One way to better understand the dimensionality of leaf trait variation is to compare models with different numbers of dimensions, and to ask if models with multiple dimensions provide improved statistical fits and conceptual in-sights compared to a single axis. For example, the two-dimensional ‘LMAm-LMAs’ model proposed by Osnas et al. (2018) could be compared to a one-dimensional LMA model in terms of their capacities to explain variation in other traits. However, implementing the ‘LMAm-LMAs’ model is challenging, because although certain leaf mass components can be neatly classified as ‘metabolic’ or ‘structural’ (Poorter et al. 2009, Osnas et al. 2018), other leaf mass components cannot. For example, thick cell walls contribute to structural toughness (Onoda et al. 2015), but at least some cell wall mass is required for the biomechanical support that enables photosynthesis. Thus, partitioning LMA into different functional components requires novel empirical or modeling approaches.

In this paper, we present a statistical modeling framework to partition LMA into metabolic and structural components: LMAm and LMAs. We develop and test this two-dimensional model using leaf trait data from two tropical forest sites (sun and shade leaves from wet and dry sites in Panama) and the GLOPNET global leaf traits database (Wright et al. 2004). We use the model to address the following questions: (1) Are measured leaf traits (including photosynthetic capacity, dark respiration rate, LL, and concentrations of nutrients and cellulose) better predicted by a one-dimensional (total LMA) or two-dimensional (LMAm-LMAs) model? (2) What are the relative contributions of LMAm and LMAs to total LMA variance in different leaf assemblages (Panama sun leaves, Panama shade leaves, and the global flora)? (3) Do LMAm and LMAs differ between evergreen and deciduous species, and between sun and shade leaves? If so, how? and (4) How do the answers to the preceding questions inform our understanding of empirical patterns of trait variation (e.g., relationships among measured leaf traits)?

## Material and Methods

### Overview

We considered multiple approaches to modeling datasets that include observations of leaf mass per area (LMA), leaf lifespan (LL), net photosynthetic capacity (*A*_max_), and dark respiration rate (*R*_dark_). These traits comprise four of the six traits in the global leaf economics spectrum (LES) analysis of Wright et al. (2004). For simplicity, we did not include the other two LES traits – leaf nitrogen (N) and phosphorus (P) concentrations – in our modeling framework. Instead, we reserved observations of leaf N and P concentrations (and cellulose content, when available) for independent model tests. We considered a simple one-dimensional model that predicts LL, *A*_max_, and *R*_dark_ from LMA alone, as well as two-dimensional models that predict LL, *A*_max_, and *R*_dark_ from two additive LMA components: structural and metabolic and structural leaf mass per area (LMAm and LMAs, respectively). We formulate the models in terms of area-normalized *A*_max_ and *R*_dark_ (*A*_area_ and *R*_area_, respectively), as these trait forms share the same denominator as LMA and its components.

To implement the two-dimensional models, we developed a statistical modeling framework to partition LMA into additive LMAm and LMAs components. We fit the models to two datasets: the GLOPNET global leaf traits dataset (Wright et al. 2004), which primarily represents interspecific variation; and the Panama dataset described by Osnas et al. (2018), which includes traits for both sun and shade leaves at wet and dry tropical forest sites. Because LMAm and LMAs are modeled (rather than observed), the two-dimensional models require one parameter per analysis unit (a species in GLOPNET or a species × canopy-position in the Panama dataset) to partition observed LMA into LMAm and LMAs. Given the large number of parameters in these models, we performed tests with randomized data to evaluate if our two-dimensional models were prone to overfitting, which could lead to spurious conclusions. These tests suggested that our two-dimensional modeling approach revealed meaningful patterns in the observed trait data.

### All statistical analyses were conducted in R version 4.2.1 (R Core Team 2022) using the R package *targets* version 0.12.0 for workflow management (Landau 2021)

### Modeling leaf lifespan, photosynthetic capacity, and dark respiration rate

We considered five types of models, ranging from simple models with LMA as the sole predictor, to more complex models in which LMA was partitioned into LMAm and LMAs. In all models, the unit of analysis is a ‘leaf sample’, defined as a species in the GLOPNET dataset or a species × canopy position in the Panama dataset (the datasets are described below).

First, we considered a simple set of models with LMA as the sole predictor for *A*_area_, *R*_area_, and LL according to power-law relationships.

Next, we considered models in which LMA is partitioned into additive metabolic and structural components for leaf sample *i*:

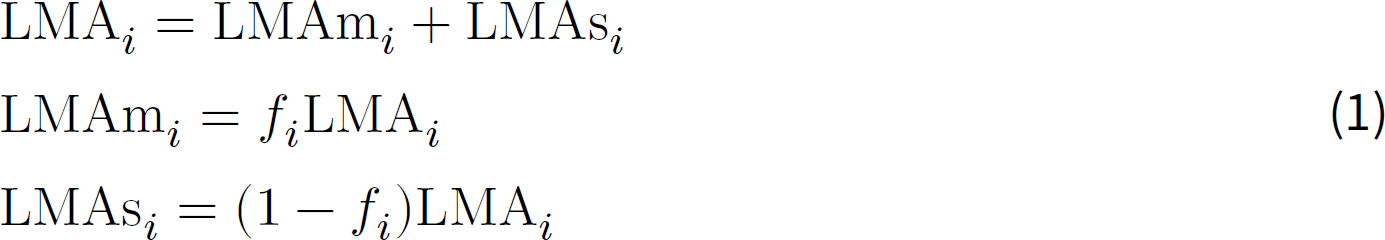

where the *f_i_* values – the fractions of LMA comprised of LMAm for each sample *i* – are estimated as latent variables in our modeling framework (see details below). We assumed that the observed values of *A*_area_, *R*_area_ and LL were related to the unobserved values LMAm and LMAs according to multivariate power-laws:

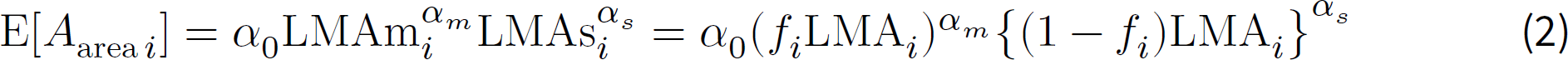

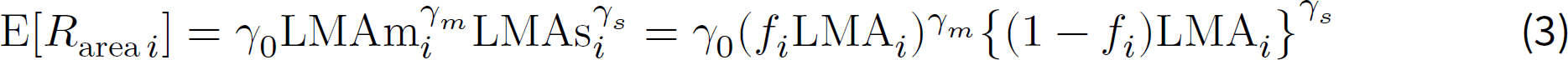

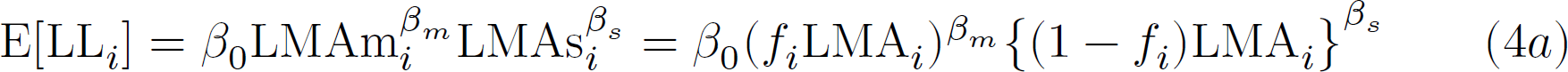

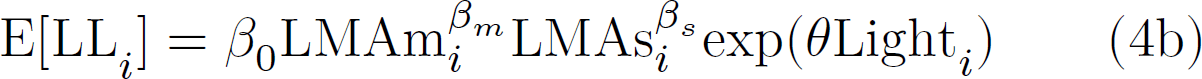

where E[⋅] indicates expected value; the ɑ, β, and γ terms are fitted parameters; and the logarithms of *A*_area_, *R*_area_, and LL are assumed to have a multivariate normal distribution (Appendix S1). In preliminary analyses, we also considered alternative (non-power-law) forms, but none of these performed better than the power-law forms and are not discussed further.

We also considered models in which LMAs in Eq. 4 was replaced by leaf structural density (LMAs/LT, where LT is leaf thickness), which is based on the observation that cellulose density is a good predictor for LL in some species assemblages (Kitajima et al. 2012, 2013). These models with LMAs/LT yielded qualitatively similar results as those based on LMAs, but did not perform as well in the model evaluation (see below).

To ensure that the model was identifiable, we imposed two broad assumptions: (i) *A*_area_ depends more strongly on metabolic leaf mass (LMAm: parameter ɑ_*m*_ in Eq. 2) than structural leaf mass (LMAs: parameter ɑ_*s*_), and (ii) LL depends more strongly on LMAs (β_*s*_ in Eq. 4 than LMAm (β_*m*_). The first assumption was implemented in different model versions either by setting ɑ_*s*_ = 0 or by imposing the constraint ɑ_*m*_ > ɑ_*s*_. Similarly, the second assumption was implemented either by setting β_*m*_ = 0 or by imposing the constraint β_*s*_ > β_*m*_. The weaker form of these assumptions (ɑ_*m*_ > ɑ_*s*_ and β_*s*_ > β_*m*_) is primarily a labeling convention and only weakly constrains the possible biological model outcomes by excluding the possibility that a single LMA component could be the primary determinant of both *A*_area_ and LL. The stronger form of the assumptions (ɑ_*s*_ = 0 and β_*m*_ = 0) leads to a more parsimonious model (fewer parameters). We considered different combinations of the strong and weak forms of the assumptions for *A*_area_ (ɑ_*m*_ and ɑ_*s*_) and LL (β_*m*_ and β_*s*_) using cross-validation (see details below). We did not impose any constraints on *R*_area_ (γ_*m*_ and γ_*s*_).

Finally, we considered a functional form that accounts for the effects of light availability on LL. This light-dependent model is motivated by optimal LL theory, which predicts decreasing LL with increasing light availability (Kikuzawa 1991), and also by the often-observed ‘LMA counter-gradient’, whereby LL and LMA positively covary across species but negatively covary within species across light gradients (Lusk et al. 2008, Russo and Kitajima 2016, Osnas et al. 2018). Mechanistically modeling how light affects LL via leaf carbon balance (Xu et al. 2017) is beyond the scope of our study. Instead, we introduced light effects in a simple way by modifying Eq. 4a to 4b where the dummy variable Light*_i_* is set to 1 for sun leaves and 0 for shade leaves, and exp(*θ*) is the sun:shade LL ratio.

### Datasets

We fit the models described above using observations of LMA (g m^−2^), *A*_area_ (mol s^−1^ m^−2^), *R*_area_ (mol s^−1^ m^−2^) and LL (months) from the GLOPNET global leaf traits database (Wright et al. 2004) and from two tropical forest sites in Panama: Parque Natural Metropolitano (PNM, “dry site”) and Bosque Protector San Lorenzo (SL, “wet site”). The GLOPNET data primarily represents interspecific variation (the dataset reports 2,548 species × site combinations, with 2,021 unique species and only 350 species occurring at more than one site) and only reports data for sun leaves if data for both sun and shade leaves are available (Wright et al. 2004). In contrast, the Panama dataset represents both inter- and intraspecific variation, including leaves sampled at two canopy positions (“sun”: full sun at the top of the canopy; and “shade”: well shaded, sampled within 2 m of the forest floor) from trees within reach of canopy cranes. The dry PNM site is a semi-deciduous coastal Pacific forest with a 5-month dry season from December-April and 1740 mm of annual rainfall (Wright et al. 2003). The PNM crane is 40 m tall with a 51 m long boom. The wet SL site is an evergreen Caribbean coastal forest with 3100 mm of annual rainfall (Wright et al. 2003). The SL crane is 52 m tall with a 54 m long boom. Additional details of the Panama dataset are described in Osnas et al. (2018).

We restricted our analysis to database records for which all four traits (LMA, *A*_area_, *R*_area_, and LL; each typically averaged over multiple leaves) were available. Each database record corresponds to an analysis unit (or ‘leaf sample’) as described above; i.e., a species in GLOPNET, or a species × canopy position in the Panama dataset. After excluding database records that lacked one or more of the four traits, 198 samples for 198 unique species were available for GLOPNET, and 130 samples for 104 unique species were available for Panama (dry and wet sites combined; 26 species sampled in both sun and shade; no species with all four traits available at both sites). In addition to LMA, *A*_area_, *R*_area_, and LL, the model based on structural leaf density also requires observations of leaf thickness, which was not available in GLOPNET but was available for 106/130 Panama samples. Both the GLOPNET and Panama datasets include additional traits that we used to interpret model results, but which were not used to fit models. These traits include nitrogen and phosphorus content per leaf unit area (*N*_area_ and *P*_area_; g m^−2^) and leaf habit(deciduous or evergreen) in both datasets, and cellulose content per unit area (CL_area_; g m^−2^) in the Panama dataset.

### Model estimation and evaluation

We modeled *A*_area_ and *R*_area_ using Eqs. 2 and 3, respectively, for both GLOPNET and Panama. To model LL for GLOPNET, we used Eq. 4a, because GLOPNET does not report canopy position (and prioritizes data for sun leaves, as described above). To model LL for Panama, we also used Eq. 4b (light effects model), which was motivated by the negative intraspecific LL-LMA relationship observed in Panama (Xu et al. 2017, Osnas et al. 2018) and elsewhere (Lusk et al. 2008, Russo and Kitajima 2016).

Posterior distributions of all parameters were estimated using the Hamiltonian Monte Carlo algorithm (HMC) implemented in Stan (Carpenter et al. 2017). We used non-informative or weakly informative prior distributions (Lemoine 2019). Prior distributions for the latent variables *f_i_*(which are used to partition LMA into LMAm and LMAs according to Eq. 1 were non-informative uniform(0, 1) distributions, (i.e., LMA was partitioned based on patterns in the data). See Appendix S1 for more detail. The Stan code use to fit models is available from Github at: https://github.com/mattocci27/LMAms. Convergence of the posterior distribution was assessed with the Gelman-Rubin statistic with a convergence threshold of 1.05 for all parameters (Gelman et al. 2013).

Because our two-dimensional (LMAm-LMAs) modeling approach includes many parameters (one latent variable *f_i_* to partition LMA into LMAm and LMAs for each leaf sample), we implemented tests with randomized data to assess potential overfitting. We generated 10 different randomized datasets by randomizing all trait values (LMA, *A*_area_, *R*_area_ and LL) across species. Thus, the randomized datasets had zero expected covariance among traits. Models fit to the randomized datasets either did not converge or showed divergent transitions (Appendix S2), indicating that indicating that models fit to randomized data do not provide reliable inferences (Betancourt 2016). Furthermore, when models were fit to randomized data, the scaling exponents (ɑ_*m,s*_, β_*m,s*_, and γ_*m,s*_) were not significantly different from zero. In simple terms, the two-dimensional models failed when fit to randomized data. In contrast, when fit to the observed (non-randomized) data, the two-dimensional models converged without divergent transitions and out-performed one-dimensional models (total LMA) for both GLOPNET and Panama (see Results). Thus, the tests with randomized data indicate that our two-dimensional approach is not inherently prone to overfitting or to creating spurious results. We therefore assume that estimates of LMAm and LMAs obtained from the GLOPNET and Panama datasets reflect meaningful patterns in the observations and allow for a meaningful exploration of our questions.

We used Pareto-smoothed importance sampling leave-one-out cross-validation (PSIS-LOO; (Vehtari et al. 2017)) to compare the performance of different models. PSIS-LOO is an accurate and reliable approximation to standard leave-one-out cross-validation (LOO), which is a robust method for comparing models with different numbers of parameters (Vehtari et al. 2017). LOO requires *n* model fits (for a dataset with *n* observations) and was therefore impractical in our study due to our computationally-demanding modeling approach. We used PSIS-LOO to calculate the LOO Information Criterion (LOOIC) for each model. A lower LOOIC indicates a better model in terms of predictive accuracy (Vehtari et al. 2017).

### Understanding relationships between photosynthetic capacity and LMA

We applied our LMAm-LMAs model to simulated data to better understand relationships between photosynthetic capacity (*A*_max_) and LMA. The causes and interpretation of relationships between *A*_max_ and LMA are controversial (Westoby et al. 2013). Although *A*_max_ is often mass-normalized (e.g., Wright et al. 2004, Shipley et al. 2006, Blonder et al. 2011), it has been argued that *A*_max_ should be area-normalized when exploring trait relationships, because photosynthesis is an area-based process (Lloyd et al. 2013). Consistent with this argument, Osnas et al. (2013) showed that across global species, variation in whole-leaf *A*_max_ is strongly dependent on leaf area, but only weakly dependent on leaf mass (after controlling for variation in leaf area). Osnas et al. (2018) further showed that the relationship between *A*_max_ and LMA (i.e., the degree of mass- vs. area-dependence) is sensitive to the amount of LL variation in an assemblage, which we hypothesize depends on the fraction of total LMA variance in the assemblage that is due to LMAs variance.

To better understand the factors affecting relationships between *A*_max_ and LMA, we created simulated datasets in which we varied the following factors: the sensitivity of *A*_area_ to variation in LMAm (parameter ɑ_*m*_ in Eq. 2), the sensitivity of *A*_area_ to LMAs (parameter ɑ_*s*_ in Eq. 2), and the fraction of total LMA variance due to variance in LMAs. For each simulated dataset, we quantified the *A*_max_ vs. LMA relationship following Osnas et al. (2018):

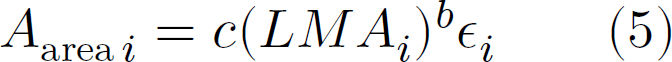

where LMA is the sum of LMAm and LMAs Eq. 1, *c* is a fitted constant, and *b* is an index of mass-dependence as illustrated by the following cases (Osnas et al. 2013, 2018): if *b* = 0, then *A*_area_ is independent of LMA, which implies that whole-leaf *A*_max_ is proportional to leaf area; conversely, if *b* = 1, then *A*_area_ is proportional to LMA, which implies that whole-leaf *A*_max_ is proportional to leaf mass. Intermediate cases (0 < *b* < 1), as well as more extreme cases (*b* ≤ 0 or *b* ≥ 1), are also possible, with reported estiamtes being between 0 and 1 (Osnas et al. 2018). Note that applying Eq. 5 to either mass- or area-normalized *A*_max_ yields equivalent results (Osnas et al. 2018).

For these simulations, we generated LMAm and LMAs values based on their estimated distributions in our GLOPNET and Panama analyses, while varying the LMAs variance (so that LMAs contributed different fractions of total LMA variance in different simulated datasets). Although Panama sun and shade leaves were analyzed together when fitting the LMAm-LMAs models Eqs. 2 - 4, they were analyzed separately here due to their different estimated covariance structures (see details below).

For GLOPNET and Panama sun leaves, LMAm and LMAs were generated independently of each other, consistent with the low estimated correlations between LMAm and LMAs in these assemblages (see Results). Specifically, LMAm values were randomly sampled from a lognormal distribution with the log-scale mean and standard deviation given by our model outputs; and LMAs values were randomly sampled from a lognormal distribution with the log-scale mean given by our model outputs and the log-scale standard deviation raging from log(1.01) to log(10) across simulated datasets. In contrast to GLOPNET and Panama sun leaves, our analysis revealed a negative correlation of *r* = −0.47 between LMAm and LMAs for Panama shade leaves (see Appendix S3). Therefore, we used a multivariate normal distribution with *r* = −0.47 to generate log(LMAm) and log(LMAs) for the Panama shade simulations. Otherwise, the simulation protocol was as described above for GLOPNET and Panama sun leaves.

For each simulated set of LMAm and LMAs values, we generated *A*_area_ values using the estimated values of ɑ_*m*_ and ɑ_*s*_ (Eq. 2) from the corresponding best-fitting model (GLOPNET or Panama); i.e., the model with the lowest LOOIC. We generated 1000 simulated datasets for GLOPNET, Panama sun leaves, and Panama shade leaves, with each dataset having a sample size of 100 leaves. For each simulated dataset, we used Eq. 5 to quantify the relationship between *A*_area_ and LMA.

### Variance partitioning

To estimate the contributions of LMAm and LMAs to total LMA variance (where LMA = LMAm + LMAs), we used the following identity:

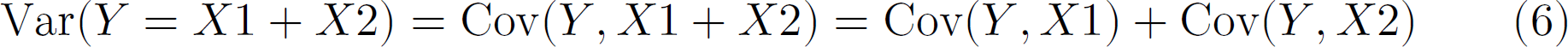

where Var(⋅) is variance and Cov(⋅) is covariance. Thus, the fractions of total LMA variance due to variance in LMAm and LMAs are Cov(LMA, LMAm)/Var(LMA) and Cov(LMA, LMAs)/Var(LMA), respectively. We applied this method to the simulated datasets described above and also to estimates of LMAm and LMAs for GLOPNET and Panama sun and shade leaves. Note that Cov(LMA, LMAs)/Var(LMA) can be greater than 100% when there is a negative covariance between LMA and LMAm.

To estimate the variation in each LMA component between and within leaf habits (evergreen vs. deciduous), sites (wet vs. dry), and light (sun vs. shade), we used ANOVA. Those post-hoc analyes were performed with posterior median parameter values.

## Results

### 1. Decomposing LMA into metabolic and structural components leads to improved pre-dictions of Aarea, Rarea and LL

For both the GLOPNET and Panama datasets, two-dimensional models (LMAm-LMAs) performed better in cross validation than one-dimensional (LMA) models (Table 1). Thus, predictions of *A*_area_, *R*_area_ and LL were improved by partitioning LMA into separate metabolic and structural components (see details below and Tables 1-2). Each of these traits (*A*_area_, *R*_area_ and LL) had a strong positive relationship with either LMAm or LMAs (but not both), which was always stronger than the corresponding relationship with total LMA (Figs. 2-3).

**Fig. 2:**
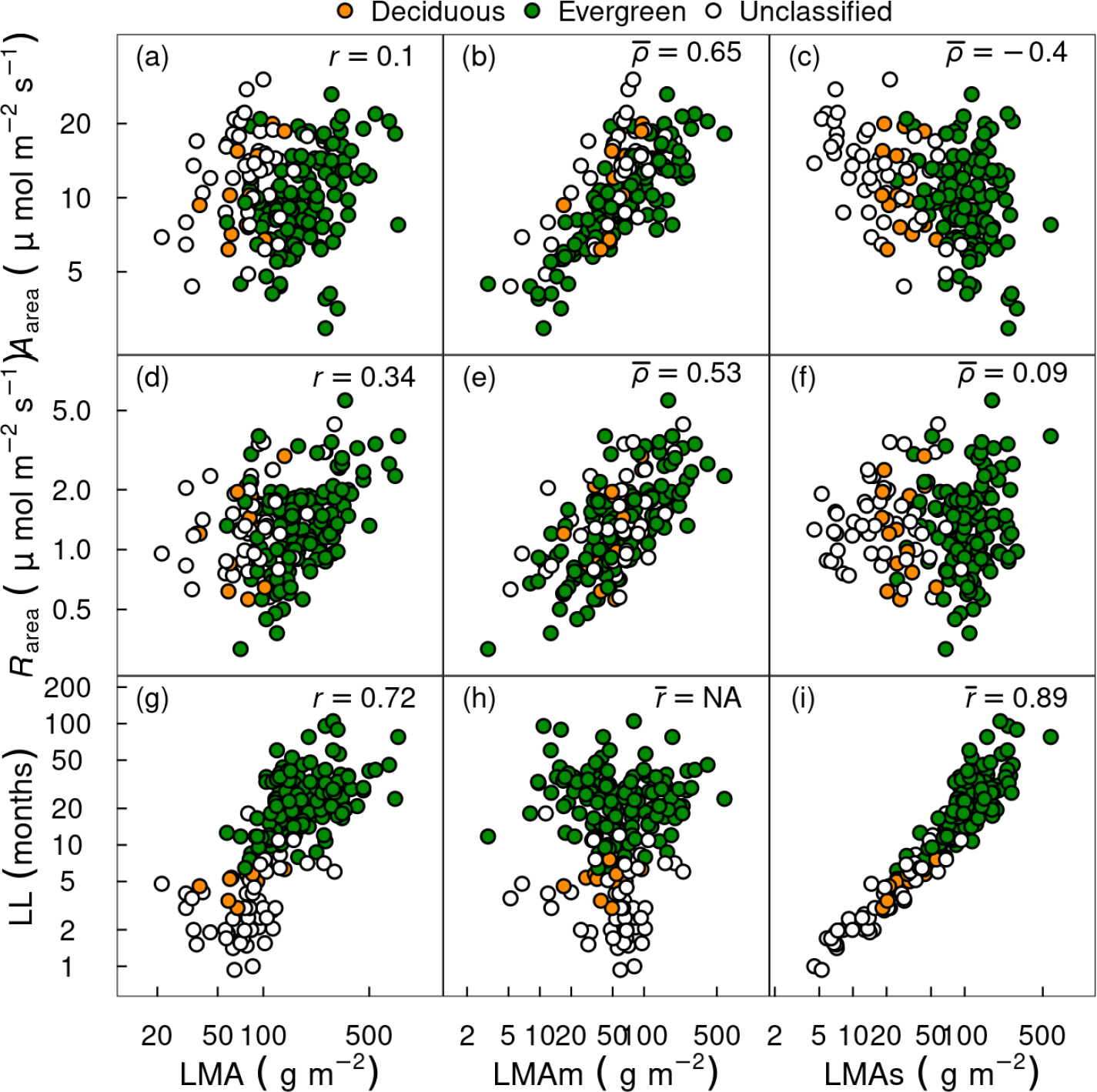
Observed and estimated leaf-trait relationships in the global GLOPNET dataset. Estimates are from the best GLOPNET model (Table 1). Leaf life span (LL), net photosynthetic rate per unit leaf area (*A*_area_), and dark respiration rate per unit leaf area (*R*_area_) are plotted against observed LMA, posterior medians of LMAm and LMAs. Pearson correlation coefficients (*r*) for LMA (left column) and posterior medians of Pearson correlation coefficients (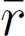) or partial correlation coefficients (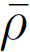) for LMAm (middle column) and LMAs (right column) are shown. Note that LL was modeled by LMAs alone due to improved model performance with this parameter constraint (Table 1). Therefore, (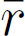) is reported for LL (single predictor variable), whereas (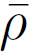) variables). is reported for *A*_area_ and *R*_area_ (multiple predictor variables).

**Table 1.**
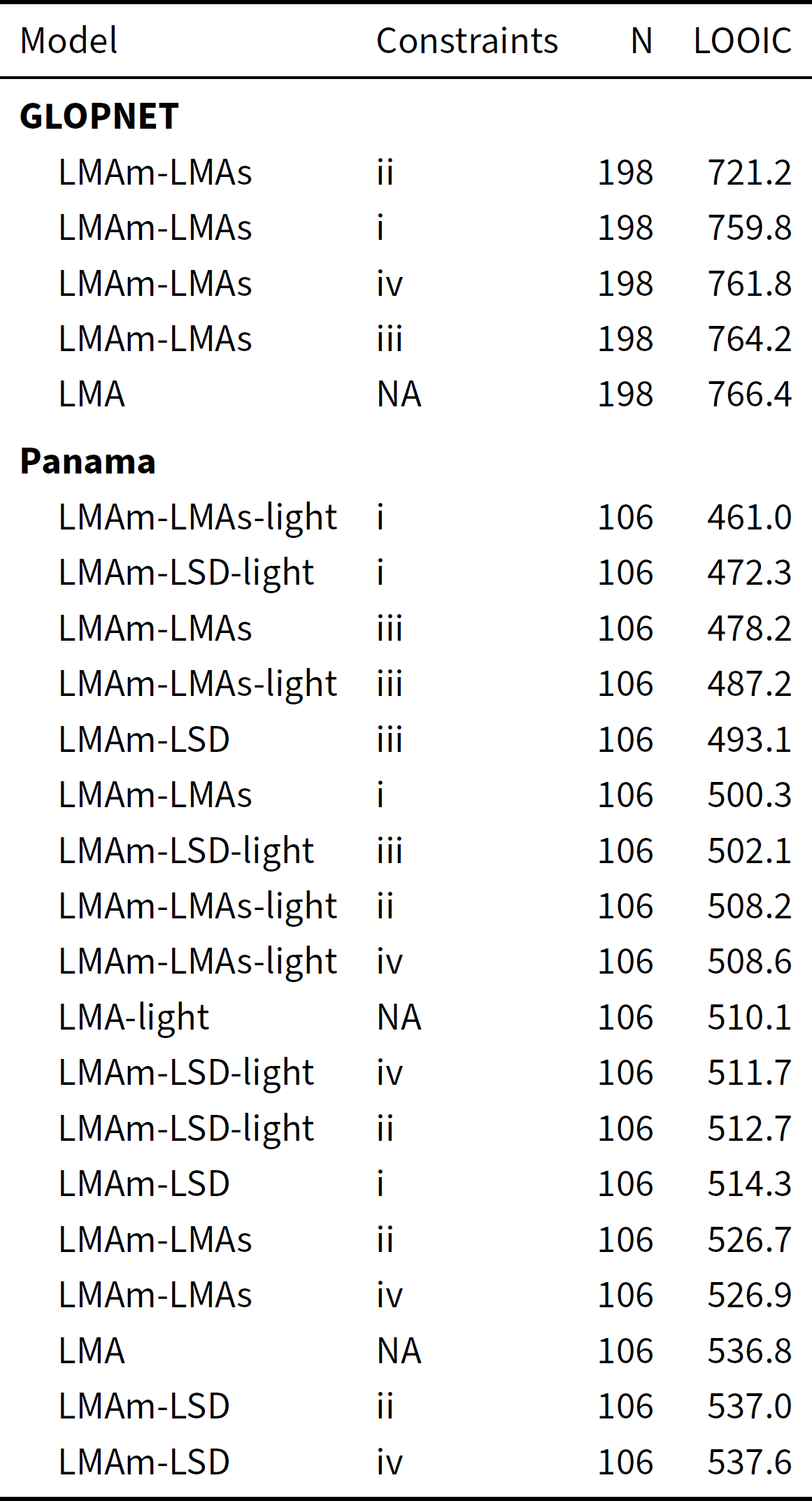
List of models fit to GLOPNET and Panama data ordered from highest to lowest cross-validation performance. Lower approximate leave-one-out cross-validation information criterion (LOOIC) indicates better predictive accuracy. For GLOPNET, we considered two model forms were considered: A_area_, R_area_, and LL were modeled as power laws of LMA (one-dimensional model) or both LMAm and LMAs (two-dimensional model; see Eqs. 2, 3, and 4a). For the Panama data, which includes canopy position (sun vs. shade) for each leaf sample, we considered the same two model forms as for GLOPNET and three additional light-dependent forms for LL (as in 4b), LMA and light (one-dimensional); LMAm, LMAs, and light (two-dimensional); and LMAm, leaf structural density (LSD = LMAs/LT), and light (two-dimensional). We combined the two-dimensional model forms with four types of parameter constraints: (i) ɑ_*s*_ = 0 and β_*m*_ = 0, (ii) β_*m*_ = 0, (iii) ɑ_*s*_ = 0, and (iv) ɑ_*m*_ > ɑ_*s*_ and β_*s*_ > β_*m*_.

| Model | Constraints | N | LOOIC |
| --- | --- | --- | --- |
| **GLOPNET** |  |  |  |
| LMAm-LMAs | ii | 198 | 721.2 |
| LMAm-LMAs | i | 198 | 759.8 |
| LMAm-LMAs | iv | 198 | 761.8 |
| LMAm-LMAs | iii | 198 | 764.2 |
| LMA | NA | 198 | 766.4 |
| **Panama** |  |  |  |
| LMAm-LMAs-light | i | 106 | 461.0 |
| LMAm-LSD-light | i | 106 | 472.3 |
| LMAm-LMAs | iii | 106 | 478.2 |
| LMAm-LMAs-light | iii | 106 | 487.2 |
| LMAm-LSD | iii | 106 | 493.1 |
| LMAm-LMAs | i | 106 | 500.3 |
| LMAm-LSD-light | iii | 106 | 502.1 |
| LMAm-LMAs-light | ii | 106 | 508.2 |
| LMAm-LMAs-light | iv | 106 | 508.6 |
| LMA-light | NA | 106 | 510.1 |
| LMAm-LSD-light | iv | 106 | 511.7 |
| LMAm-LSD-light | ii | 106 | 512.7 |
| LMAm-LSD | i | 106 | 514.3 |
| LMAm-LMAs | ii | 106 | 526.7 |
| LMAm-LMAs | iv | 106 | 526.9 |
| LMA | NA | 106 | 536.8 |
| LMAm-LSD | ii | 106 | 537.0 |
| LMAm-LSD | iv | 106 | 537.6 |

**Table 2.** Posterior medians [95% credible interval] of parameters for the best GLOPNET and Panama models. Bold values are significantly different from zero based on the 95% credible interval.

| Parameters | GLOPNET | Panama |
| --- | --- | --- |
| Effect of LMAm on $A_{\text{area}} (\alpha_p)$ | **0.281 [0.204, 0.36]** | **0.577 [0.429, 0.723]** |
| Effect of LMAs on $A_{\text{area}} (\alpha_s)$ | **-0.131 [-0.201, -0.058]** | - |
| Effect of LMAs on LL ($\beta_s$) | **0.887 [0.761, 1.018]** | **0.352 [0.17, 0.531]** |
| Effect of LMAm on $R_{\text{area}} (\gamma_p)$ | **0.289 [0.202, 0.38]** | **0.614 [0.388, 0.823]** |
| Effect of LMAs on $R_{\text{area}} (\gamma_s)$ | 0.035 [-0.045, 0.115] | 0.032 [-0.25, 0.351] |
| Effect of light on LL ($\theta$) | - | **-0.83 [-1.116, -0.538]** |

In the best GLOPNET model (lowest LOOIC), *A*_area_ increased with LMAm and decreased with LMAs, *R*_area_ increased with LMAm and had little dependence on LMAs, and LL increased with LMAs and was independent of LMAm (Tables 1-2; Fig. 2).

In the best Panama model, *A*_area_ increased with LMAm and was independent of LMAs, *R*_area_ increased with LMAm and had little dependence on LMAs, and LL increased with LMAs and was independent of LMAm (Tables 1-2; Fig. 3). The best Panama model also included the effect of light (sun vs. shade) on LL (Tables 1-2; Fig. 4). The light effect was not tested for GLOPNET because GLOPNET does not report canopy position.

**Fig. 3:**
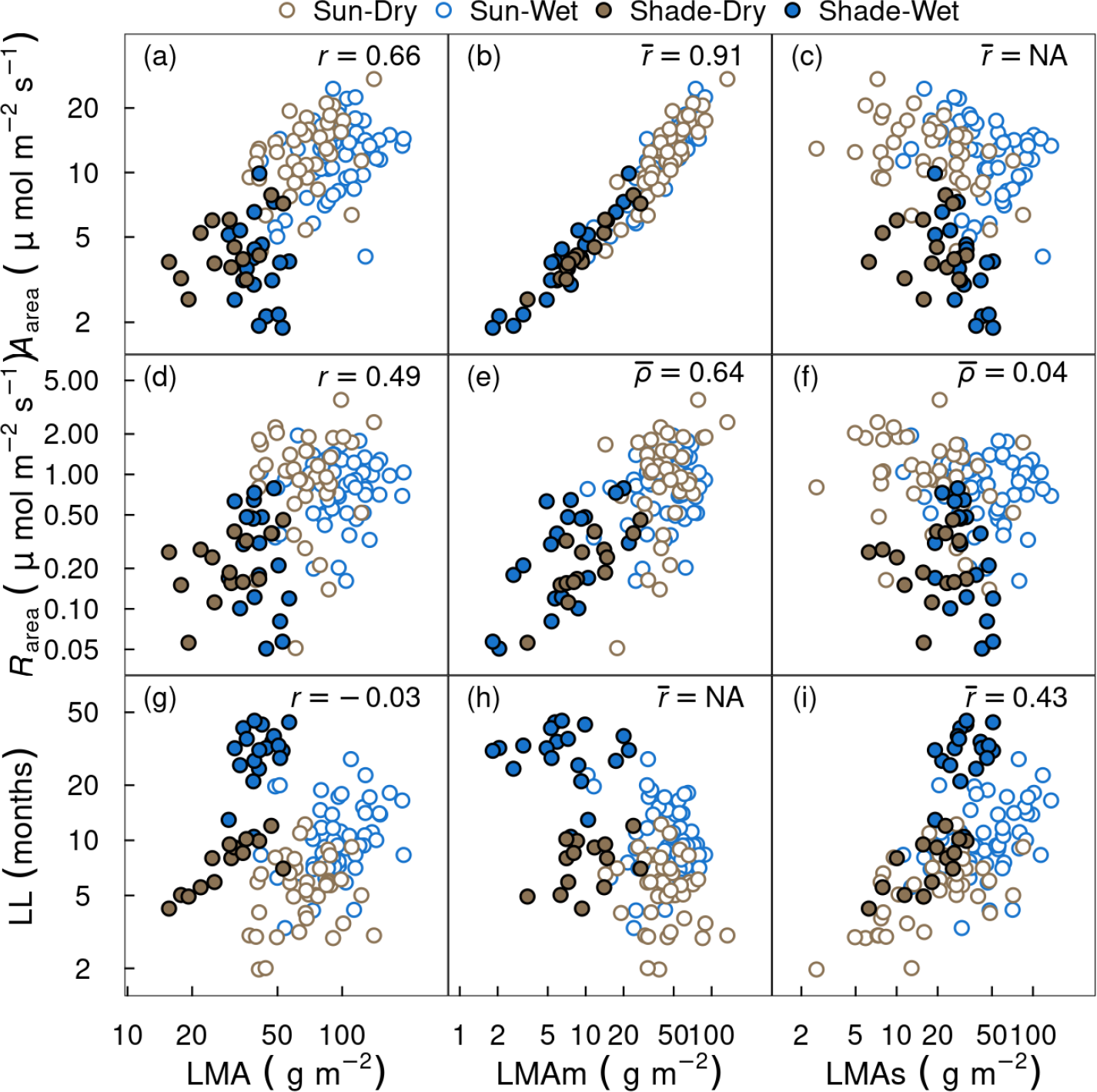
Observed and estimated leaf-trait relationships in the Panama dataset. Estimates are from the best Panama model (Table 1), which included effects of light on LL. Details as for Fig. 2. Note that *A*_area_ and LL were modeled by LMAm and LMAs alone, respectively, due to improved model performance with these parameter constraints (Table 1). The results shown here include all leaves in the Panama dataset. The observed separation of LL between sun and shade leaves is accounted for in the model predictions Fig. 4.

**Fig. 4:**
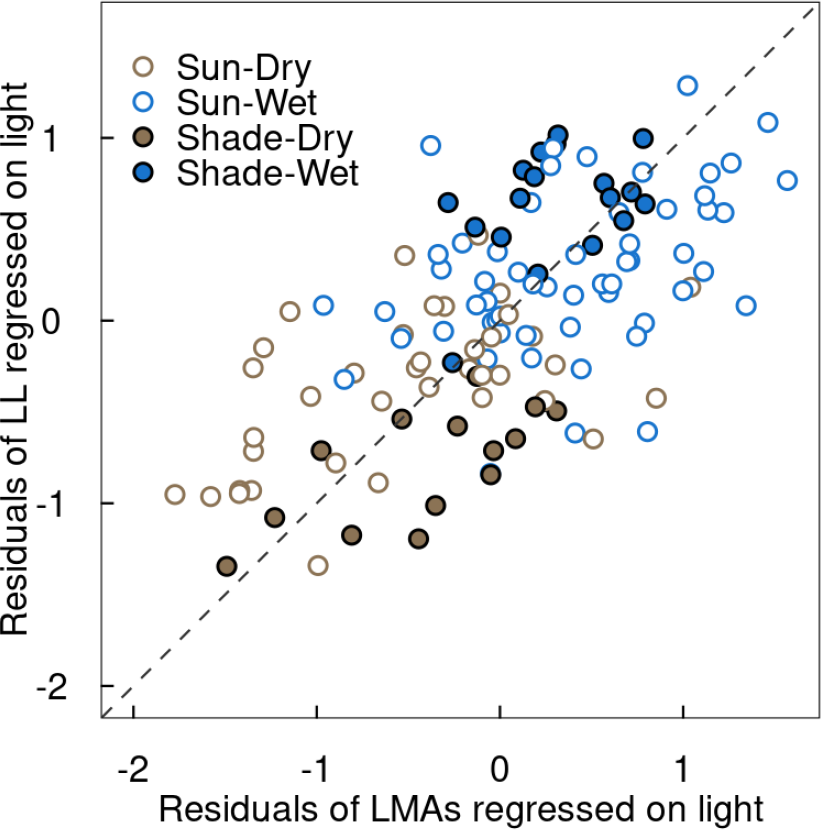
Relationship between leaf lifespan (LL) and LMAs in the Panama dataset, after accounting for the effects of light (suv vs. shade leaves; see Eq. 4b). The dashed line indicates the 1:1 relationship expected for residuals on the log-scale. The posterior medians of the partial correlation coefficient (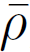) the Panama dataset. is shown. The results shown here include all leaves in the Panama dataset.

### 2. Nearly all leaf dark respiration is associated with metabolic leaf tissue mass

According to our model results, metabolic leaf mass (LMAm) accounted for nearly all leaf dark respiration; i.e., estimated dark respiration rate per-unit structural mass (γ_*s*_) was close to zero for both GLOPNET and Panama (Table 2). Thus, although building costs appear similar for different leaf chemical components and tissues (Williams et al. 1989, Villar and Merino 2001), our results suggest that leaf mass associated with structural function (i.e., toughness and LL) has roughly zero maintenance cost.

### 3. Evergreen leaves have greater LMAs than deciduous leaves, and sun leaves have both greater LMAm and LMAs than shade leaves

Leaf habit (evergreen vs. deciduous) explained very little variation in LMAm but substantial variation in LMAs for both GLOPNET and Panama (Fig. 5a-b). Thus, the higher total LMA in evergreen compared to deciduous leaves was due to differences in LMAs (Figs. 6, S1, and S2). In the Panama dataset, light (sun vs. shade) explained much more variation in LMAm than LMAs, whereas site (wet vs. dry) explained much more variation in LMAs than LMAm (Fig. 5c). Panama wet-site leaves had higher total LMA than dry-site leaves due to greater LMAs (Figs. 7 and S1). In contrast, Panama sun leaves had higher total LMA than shade leaves primarily due to greater LMAm, with a smaller but significant contribution from LMAs (Figs. 7 and S1). Thus, LMAs comprised a larger fraction of total LMA for shade leaves than for sun leaves (Fig. S2).

**Fig. 5:**
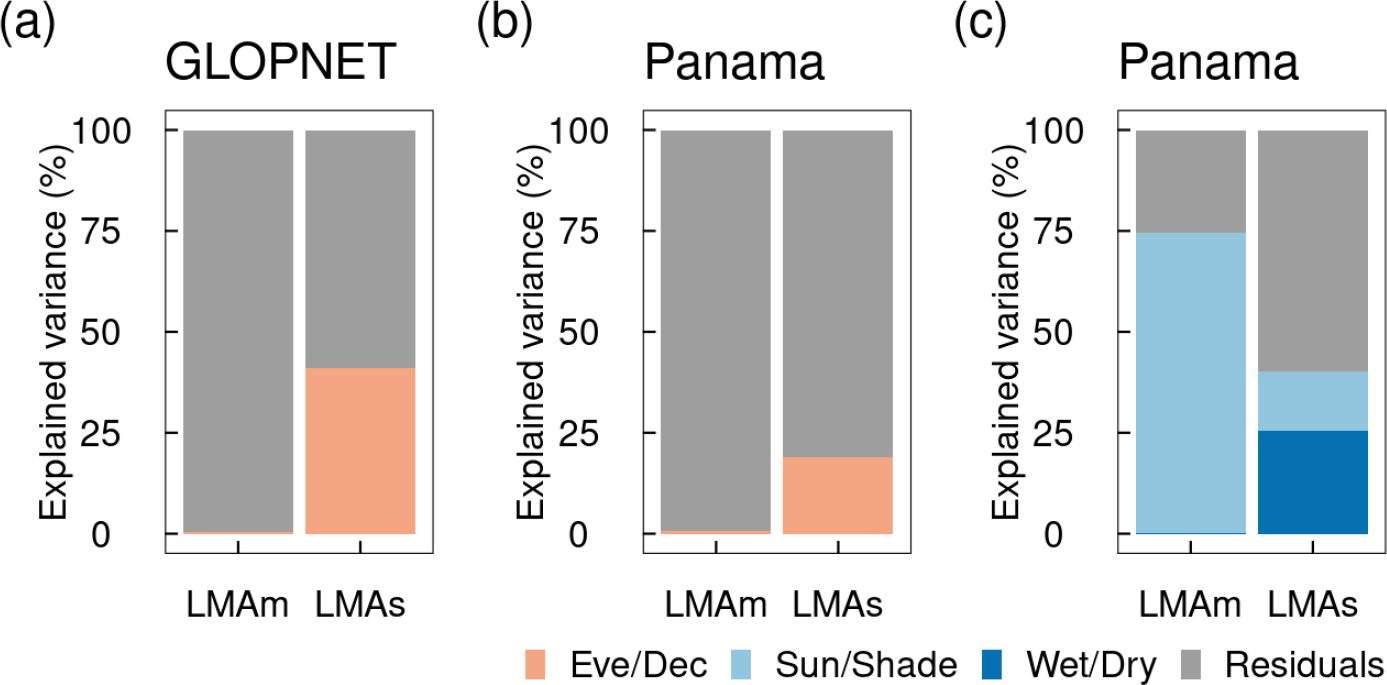
Variance partitioning on LMA components between and within leaf habits (evergreen vs. deciduous) for the GLOPNET dataset and the Panama dataset, and between and within sites (wet vs. dry) and light (sun vs. shade) for the Panama dataset. To isolate the effects of intraspecific variation, the Panama results shown here only include species for which both sun and shade leaves were available.

**Fig. 6:**
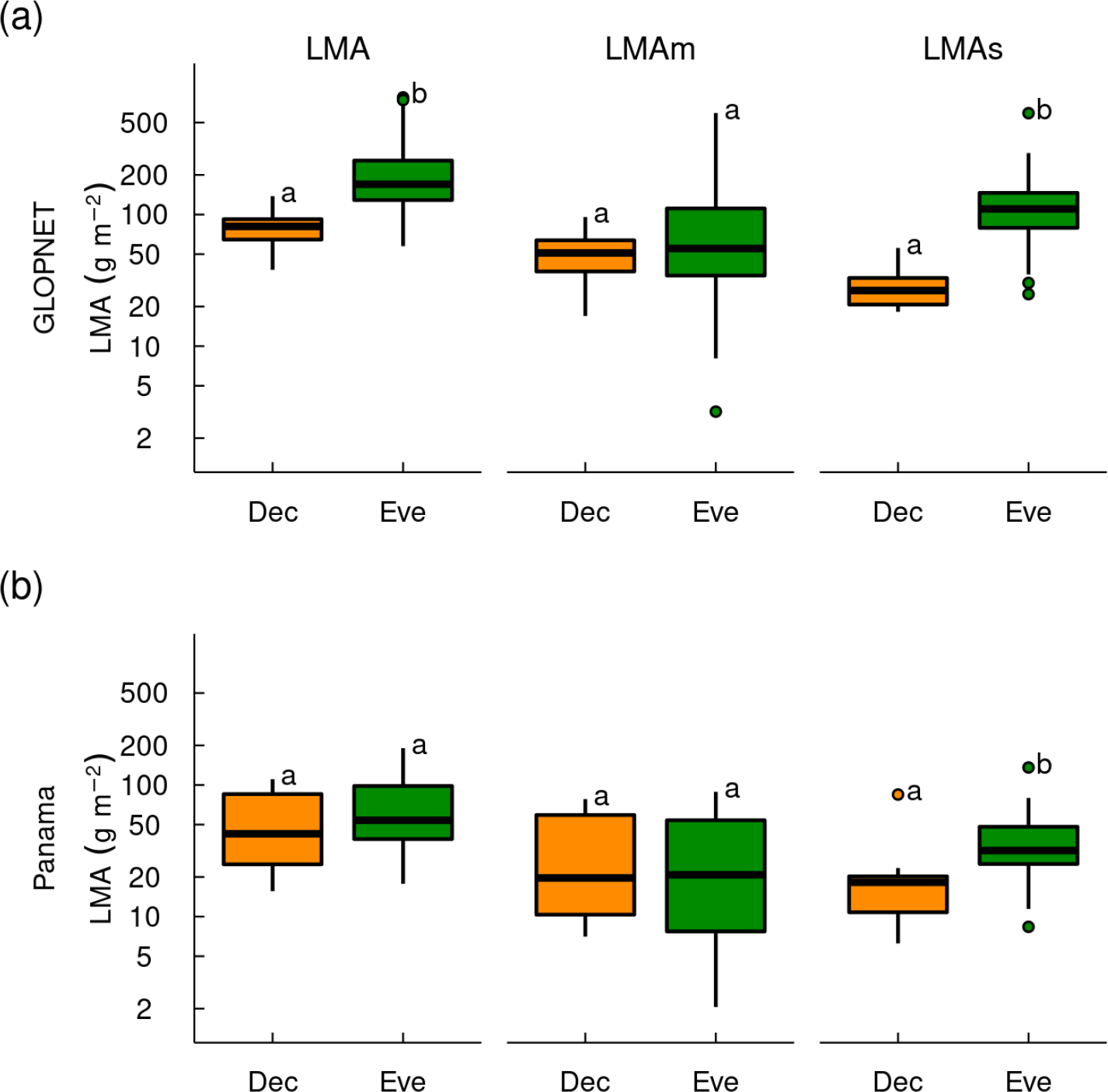
Boxplots comparing leaf mass per area (LMA) and posterior medians of photosynthetic and structural LMA components (LMAm and LMAs, respectively) across deciduous (Dev) and evergreen (Eve) leaves in the GLOPNET dataset (a) and in the Panama dataset (b). The center line in each box indicates the median, upper and lower box edges indicate the interquartile range, whiskers show 1.5 times the interquartile range, and points are outliers. Groups sharing the same letters are not significantly different (P > 0.05; t-tests). The Panama results only include species for which both sun and shade leaves were available. Qualitatively similar results were obtained when all Panama species were included (Fig. S1). Note that the vertical axis is on a log_10_ scale.

**Fig. 7:**
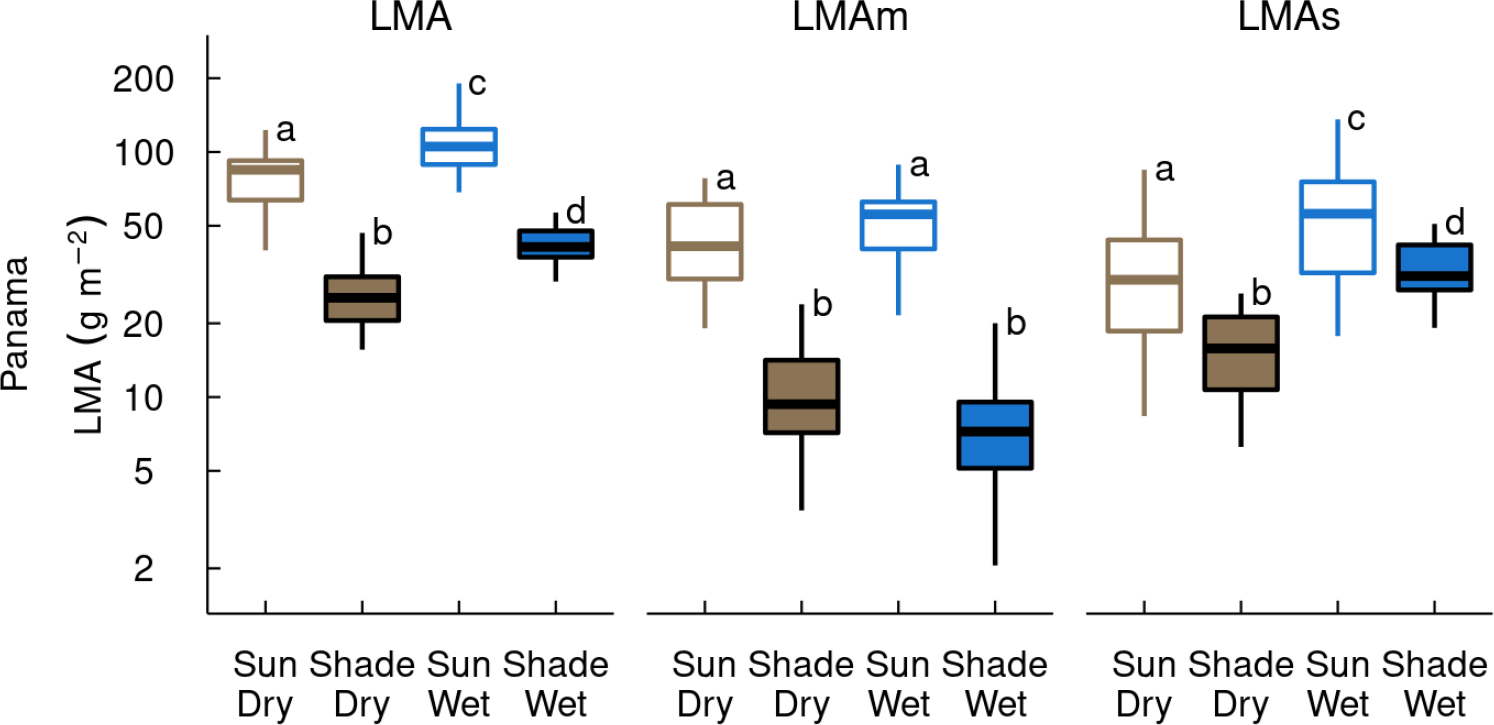
Boxplots comparing leaf mass per area (LMA) and posterior medians of metabolic and structural LMA components (LMAm and LMAs, respectively) across sites (wet and dry) and canopy strata (sun and shade) in the Panama dataset. These results only include species for which both sun and shade leaves were available. Qualitatively similar results were obtained when all Panama species were included (Fig. S1). Details as for Fig. 6.

### 4. LMA variance components affect the area- vs. mass-dependence of leaf photosynthetic capacity (Amax)

Analysis of simulated data showed that the percent of LMA variation due to LMAs variation (Eq. 6) strongly affected the area- vs. mass-dependence of *A*_max_ (i.e., the degree to which whole-leaf *A*_max_ depends on leaf area or mass; Osnas et al. (2013); Osnas et al. (2018)). Generally, mass-dependence of *A*_max_ decreased (area-dependence increased) with increasing LMAs variance (Fig. 8). Other factors, including the sensitivity of *A*_max_ per-unit leaf area (*A*_area_) to variation in LMAm and LMAs (ɑ_*m*_ and ɑ_*s*_ in Eq. 2) and covariance between LMAm and LMAs, also affect the mass-dependence of *A*_max_ (Fig. 8, Figs. S4-S5). For example, although LMAs variance was relatively low in GLOPNET (51% of total LMA variance), variation in *A*_max_ in GLOPNET was strongly area-dependent because the relatively weak dependence of *A*_area_ on LMAm and the negative dependence of *A*_area_ on LMAs (Table 2) led to weak overall dependence of *A*_area_ on LMA. Thus, mass-dependence of *A*_max_ in GLOPNET was similar or weaker than in the Panama data, despite LMAs variance comprising a larger fraction of LMA variance in Panama (74% and 100% of total LMA variance for Panama sun and shade leaves, respectively).

**Fig. 8:**
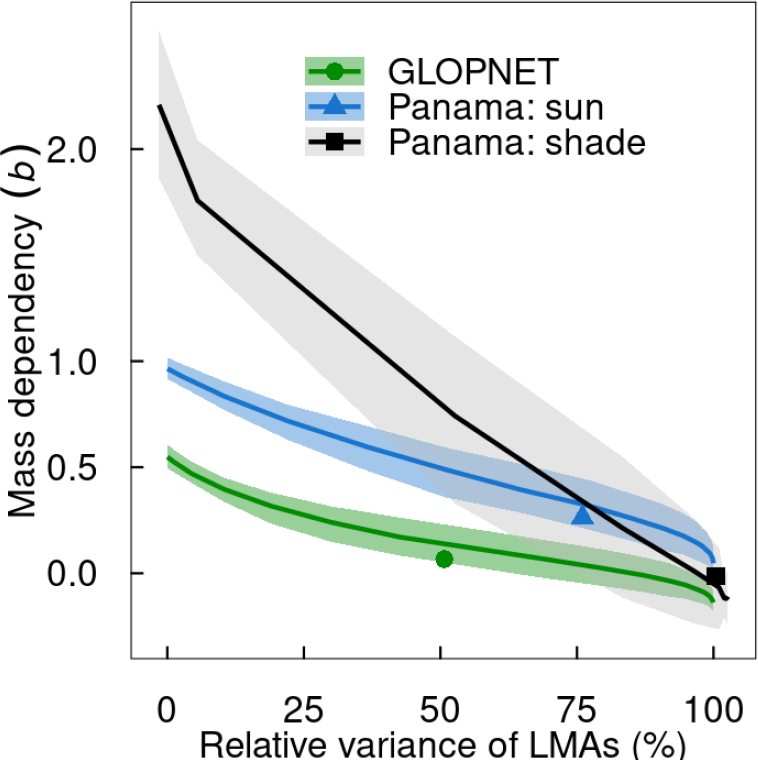
Relationships between mass dependency (*b* in Eq. 5) and LMAs variance (relative to total LMA variance; Eq. 6) for the global GLPNET dataset, sun leaves in Panama, and shade leaves in Panama. Solid lines indicate simulated medians and shaded regions indicate 95% confidence intervals. Each point indicates the estimated values from the empirical data and represents interspecific variation (e.g., across species within a canopy position in the Panama dataset). The y-axis values indicates if photosynthetic capacity (*A*_max_) is is primarily mass-dependent (*b* > 0.5) or primarily area-dependent (0.5 > *b* > 0), with *b* near 0 indicating purely area-dependendence (Osnas et al. 2018). If *b* > 1, then *A*_max_ increases exponentially with LMA, which is not consistent with observed relationships (Osnas et al. 2018).

### 5. Nitrogen and phosphorus per-unit leaf area are strongly correlated with LMAm, and cellulose per-unit leaf area is strongly correlated with total LMA

In the GLOPNET dataset, *N*_area_ and *P*_area_ had strong positive correlations with LMAm, but only weak correlations with LMAs (Fig. S6). Similarly, in the Panama dataset, *N*_area_ and *P*_area_ had strong positive correlations with LMAm but not with LMAs (Fig. 9). Cellulose content per-unit leaf area (CL_area_), which was only available for the Panama dataset, was most strongly correlated with total LMA, but also showed a clear relationship with LMAs that was consistent across canopy positions and sites (Fig. 9). Partial regression plots showed that the higher *N*_area_, *P*_area_, and CL_area_ in sun leaves (compared to shade leaves; Fig. 9) was due to sun leaves having higher LMAm than shade leaves (Fig. S7).

**Fig. 9:**
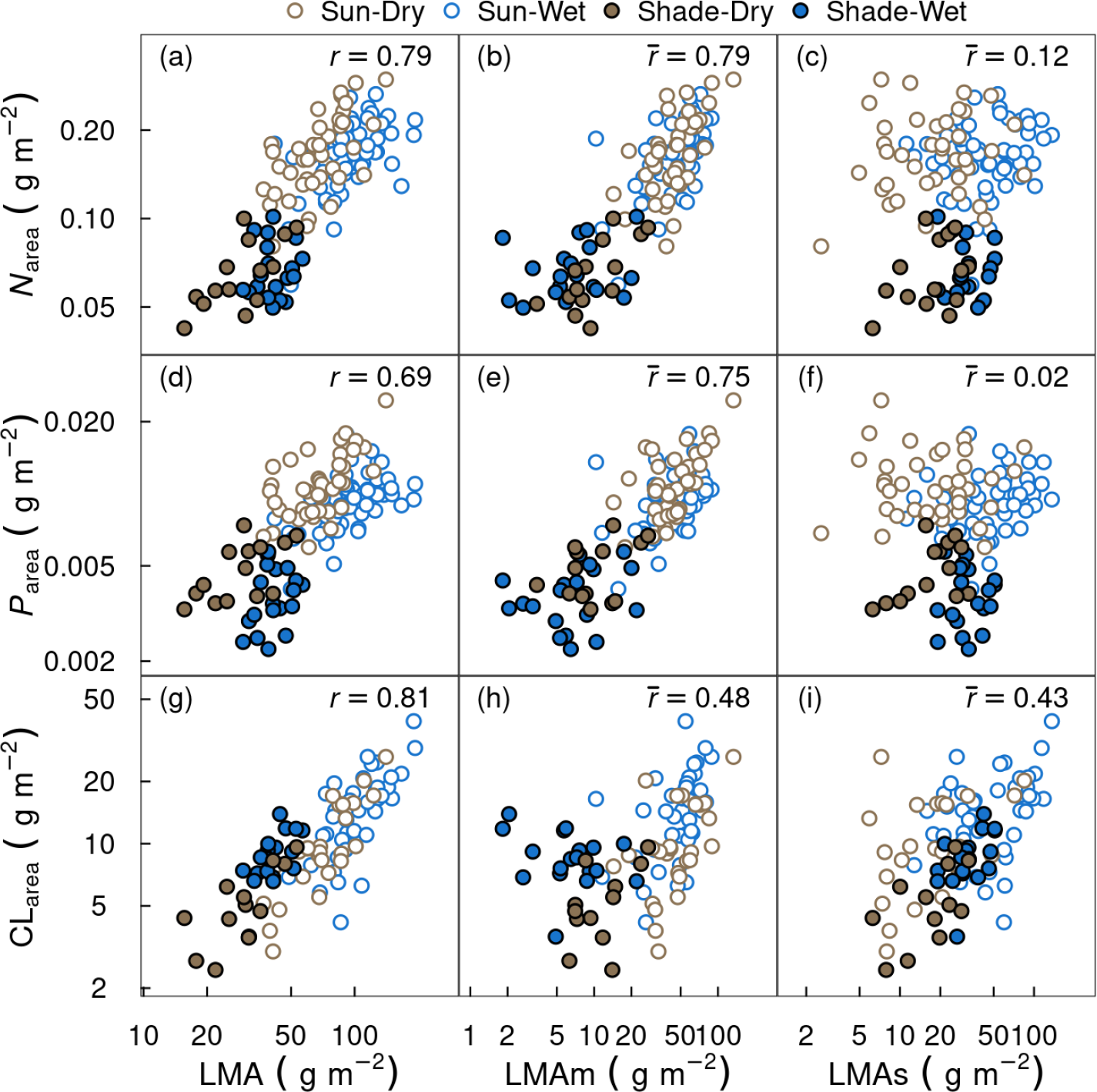
Measured traits in the Panama dataset related to photosynthesis and metabolism (nitrogen and phosphorus per-unit leaf area; *N*_area_ and *P*_area_) are better correlated with estimates (posterior medians) of the metabolic LMA component (LMAm) than the structural component (LMAs), whereas the opposite pattern occurs for a measured structural trait (cellulose per-unit leaf area; CL_area_). *N*_area_, *P*_area_, and CL_area_ data were not used to fit the models, and are presented here as independent support for the model results. Pearson correlation coefficients (*r*) for LMA and the posterior medians of Pearson correlation co-efficients (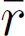) for LMAm and LMAs are shown. Analogous results were obtained for *N*_area_ and *P*_area_ for GLOPNET (Fig. S6). The results shown here include all leaves for which *N*_area_, *P*_area_ and CL_area_ are available.

## Discussion

Our analysis of the GLOPNET global dataset and a dataset including sun and shade leaves from wet and dry Panama sites demonstrates that decomposing LMA variation into separate metabolic and structural components (LMAm and LMAs, respectively) leads to improved predictions of photosynthetic capacity (*A*_max_), dark respiration rate (*R*_dark_), and leaf lifespan (LL) relative to a simpler one-dimensional model based on total LMA. Furthermore, LMAm and LMAs showed clear relationships with independent model evaluation data (nitrogen, phosphorus, and cellulose content per-unit leaf area). In our analysis, LMAm and LMAs are model-inferred (rather than observed) traits. Nevertheless, several observations suggest that we can derive meaningful insights from these modeled LMA components. Firstly, our analysis revealed several patterns that are consistent with previous analyses of measured traits (summarized below). Secondly, although our modeling approach requires many parameters to partition LMA into LMAm and LMAs (one latent variable per leaf sample), which could lead to over-fitting or spurious results, our model performed poorly (often failing to converge) when applied to randomized data. Thus, the model structure is not prone to creating signals from noise. Together, these observations lend credibility to our model-inferred estimates of LMAm and LMAs. We first summarize and interpret our model results in the context of previous studies. We then discuss the new insights derived from our analysis and implications for vegetation models, including the land components of Earth system models.

### Modeling *A*_max_, *R*_dark_, and LL as functions of metabolic and structural LMA

*A*_max_ per-unit leaf area (*A*_area_) increased with LMAm and was either unrelated to LMAs (Panama) or decreased with LMAs (GLOPNET; Figs. 2-3 and Table 2). The decline in GLOPNET *A*_area_ with increasing LMAs may be due to decreasing mesophyll conductance, which decreases with increasing cell wall thickness (Evans et al. 2009, Terashima et al. 2011, Onoda et al. 2017). This trend may be less apparent in our Panama analysis due to the relatively small LMA range (16-193 g m^2^ in Panama, compared to 21-776 g m^2^ in GLOPNET) or may be obscured by pooling sun and shade leaves in a single analysis. In any case, an increase in *A*_area_ with LMAm but not LMAs, as observed for both GLOPNET and Panama, is consistent with the view that LMA is a composite trait comprised of metabolically active and inactive components (Poorter et al. 2009, Osnas et al. 2018, Lichstein et al. 2021).

*R*_dark_ per-unit leaf area (*R*_area_) increased with LMAm and was unrelated to LMAs (Figs. 2-3 and Table 2), consistent with the view that LMAs is comprised of non-metabolic leaf mass components, such as cellulose, lignin, and other fibres (Onoda et al. 2011, Kitajima et al. 2012, 2016). Median values of LMAs, as a percent of total LMA, were ∼60% for evergreen species and ∼40% for deciduous species (Fig. S2), and 15-40% for shade leaves and 50-70% for sun leaves (Fig. S3). These estimates align with measurements of cell wall (fibre) content from a global dataset (14–77%, median 47% of total leaf mass, with higher values for shade than sun leaves; Onoda et al. 2011) and a Panama study site (Barro Colorado Island, BCI: ∼50% mean across species, with higher values for shade leaves; Kitajima et al. 2016).

LL increased with LMAs but was unrelated to LMAm (Figs. 2-3 and Tables 1-2), consistent with well-established relationships between leaf fibre content, toughness, and LL (Onoda et al. 2011, 2017, Kitajima et al. 2012, 2016). For a given LMAs, shade leaves had longer LL than sun leaves (see Fig. 3i and parameter *θ* in Table 2). Thus, partitioning LMA into LMAm and LMAs is not sufficient to explain the widely observed counter-gradient, whereby LL increases with increasing LMA across species but decreases with increasing LMA within species from shade to sun (Lusk et al. 2008, Russo and Kitajima 2016, Osnas et al. 2018). The failure of LMAs to provide robust predictions of LL across canopy positions may reflect factors unlreated to leaf toughness - e.g., how optimal LL depends on light availability (Kikuzawa 1991) - as well as variation in leaf toughness for a given value of LMAs. In a global meta-analysis, leaf toughness was strongly related to toughness per-unit leaf tissue density, but only weakly related to leaf thickness or density (Onoda et al. 2011). Accordingly, on BCI, Panama, the difference between shade and sun leaves was greater for cellulose content (% of leaf mass) than cell wall content (which includes cellulose, as well as other fibres that provide less structural toughness). Thus, just as not all LMA is equal, it is almost certainly true that not all LMAs is equal, with some LMAs components (e.g., cellulose) imparting greater toughness and longevity than others (e.g., lignin in secondary cell walls of vascular tissue, Kitajima et al. 2016).

In summary, the dependencies of *A*_area_, *R*_area_, and LL on model-inferred values of LMAm and LMAs are broadly consistent with previous analyses of measured traits. Our results support the hypothesis that LMA can be partitioned into metabolic and structural components, the latter having a maintenance respiration rate indistinguishable from zero. Like total LMA, LMAs is likely a composite trait, including cuticle and fibres with different functions. A more refined modeling approach that links different structural components to specific functions (e.g., mesophyll cell wall thickness that imparts overall lamina toughness vs. fibres in vascular tissue that contribute to cavitation resistance) could potentially provide a mechanistic explanation for LL variation across canopy positions as well as ecophysiological tradeoffs across edaphic-climate gradients.

### Implications for understanding LL variation

Optimal LL theory (Kikuzawa 1991) predicts that LL should increase with toughness (which increases building costs and decreases the rate of photosynthetic decline with leaf age) and decrease with initial (pre-aging) net assimilation rate. Consistent with these predictions, our results indicate an increase in LL with increasing LMAs and decreasing light. However, our results do not support a decrease in LL with increasing LMAm (which, along with the environmental conditions, should determine net assimilation rate). For example, within each Panama site (wet and dry), *A*_area_ of sun leaves varied by over a factor of four and was strongly determined by LMAm (Fig. 3b). Yet, the best-performing models excluded effects of LMAm on LL (Table 1). Modeling LL as a function of LMAs, rather than total LMA, partially explains the sun-shade LL difference (compare Figs. 3g and 3i), and explicit consideration of fibre content and toughness may provide a more complete understanding of LL variation with light (see details above). Thus, while optimal LL theory provides important insights regarding the coordination of structural toughness and net assimilation rate, toughness itself may provide a sufficient proximate explanation for LL variation.

### Implications for understanding relationships between photosynthetic capacity and LMA

Osnas et al. (2018) suggested that in leaf assemblages where LMA variation is largely due to variation in structural leaf mass components that contribute to toughness but not photosynthetic capacity, metabolic traits (including *A*_max_) should be primarily area-dependent rather than mass-dependent. Our simulations show that all else being equal, area-dependence of *A*_max_ increases (mass-dependence decreases) as LMAs variance increases (Fig. 8), as hypothesized by Osnas et al. (2018). However, our analysis also revealed complexities that were not anticipated by Osnas et al. (2018). For example, if *A*_area_ only weakly increases with LMAm and/or decreases with LMAs (as in GLOPNET; Table 2), then *A*_max_ can be strongly area-dependent (weakly mass-dependent) even if LMAs accounts for relatively little LMA variance (Fig. 8). We conclude that area- vs. mass-dependence depends not only LMAs variance, but also on how metabolic traits vary with different LMA components.

Several factors could contribute to the less-than-proportional increase in *A*_area_ with LMAm (ɑ_*p*_ < 1 for both GLOPNET and Panama; Table 2). High LMAm may lead to low mesophyll conductance, e.g., due to a longer CO_2_ diffusion pathway (Evans et al. 2009). Similarly, high LMAm may lead to a high level of chloroplast self-shading within a leaf (Terashima et al. 2011). Both of these factors would decrease the marginal benefit of additional units of LMAm, and this marginal decrease may be especially strong in GLOPNET due to its large LMAm range (3.2 - 591 g m^2^) relative to Panama (3.2 - 591 g m^2^).

Finally, while much of the model-inferred LMAm may be directly or indirectly related to photosynthesis, ‘LMAm’ may also include metabolic leaf mass unrelated to photosynthesis and/or non-metabolic leaf mass that is statistically assigned (by default) to LMAm simply because it does not affect LL. For example, non-structural carbohydrates, which are ignored in our model, can account for 5-8% of leaf mass in evergreen species (Würth et al. 2005) and up to 20% of leaf mass in evergreen species (Martínez-Vilalta et al. 2016). A more refined model that accounts for more than two LMA components may reveal stronger relationships between metabolic traits (*A*_area_ and *R*_area_) and LMAm.

In addition to providing insights as to why metabolic traits tend to be area-dependent across species, our analysis also helps explain why these same traits tend to be mass-dependent within species across light levels (Osnas et al. 2018). LMA increased from shade to sun primarily due to an increase in LMAm (Figs. 7 and S1), which should lead to strong mass-dependence of *A*_max_ and *R*_dark_ (Fig. 8) and likely reflects an increase in the size and number of palisade mesophyll cells with increasing light levels (Onoda et al. 2008, Terashima et al. 2011). Estimates of mass-dependence of metabolic traits (parameter *b* in Eq. 5, with *b* > 0.5 indicating greater mass-than area-dependence) across light levels are around 0.8-1.0 for the Panama dataset we analyzed (Osnas et al. 2018). Although values of *b* greater than 1 are theoretically possible (which implies an exponential increase in area-normalized traits with increasing LMA), two factors likely constrain the observed values of *b*. Firstly, at least some of the LMA increase from shade to sun is due to LMAs (Figs. 7 and S1), which may reflect the additional LMAs needed to protect and support the high LMAm of sun leaves and/or additional LMAs required to withstand hydraulic or other stress in the upper canopy. Secondly, as discussed above, *A*_area_ and *R*_area_ increase at a less-than-proportional rate with increasing LMAm, which in turn moderates the LMA scaling exponent (*b*).

### Implications for terrestrial ecosystem models

For over two decades, low-dimensional trait tradeoffs have been explored as a means to represent plant functional diversity in the vegetation components of biogeochemical models (Moorcroft et al. 2001, Bonan et al. 2002, Verheijen et al. 2013), with the ultimate goal of adding ecological and physiological realism to coupled Earth System Models (ESMs, Moorcroft 2006, Wullschleger et al. 2014). Recently, there has been a proliferation of interest in this topic (Scheiter et al. 2013, Sakschewski et al. 2015, Fisher et al. 2015, Sakschewski et al. 2016, Dantas de Paula et al. 2021) along with demographic approaches capable of representing the dynamics of diverse communities in ESMs (Fisher et al. 2018). The leaf economics spectrum (LES) has been interpreted as a single dominant axis of global leaf functional variation (Wright et al. 2004), and this simple framework has been adopted in some of these dynamic vegetation models. Our results demonstrate the value of considering metabolic and structural leaf mass separately, and indicate weak if any interspecific correlation between these LMA components (Fig. S8). We therefore conclude that a one-dimensional representation of leaf functional variation is too simplistic. Consistent with our conclusion that leaf function is multi-dimensional, analyses of leaf nitrogen indicate significant independent contributions of metabolic (e.g., rubisco) and structural leaf nitrogen components (Dong et al. 2017, 2022), with over 20% of total leaf nitrogen in high-LMA leaves occurring in cell walls (Onoda et al. 2017). Similarly, partitioning leaf phosphorus reveals substantial, LMA-dependent metabolic, structural, and nucleic acid fractions (Hidaka and Kitayama 2011). We suggest that future biogeochemical model developments consider multiple axes of leaf functional diversity, and that models with coupled carbon-nutrient cycles consider how LMA, nitrogen, and phosphorus are allocated to different functions

## Acknowledgments

We thank Jonathan Dushoff for statistical advice and Martijn Slot for helpful comments that improved the paper. Mirna Samaniego and Milton Garcia provided indispensable assistance in data collection. We thank the Andrew W. Mellon Foundation for supporting this work. MK was supported by a Xishuangbanna State Rainforest Talent Support Program, a CAS President’s International Fellowship Initiative (2020FYB0003), a ZiHui (Wisdom) Yunnan Program (202203AM140026), and a Postdoctoral Fellowship for Research Abroad from the Japan Society for the Promotion of Science (25-504).

## Author contributions

MK, JWL and JLDO conceived of the study; KK provided guidance on leaf physiology and interpreting trait data; KK, SJW and SAVB contributed data; MK devised the analytical approach and performed analyses; MK and JWL wrote the first draft of the manuscript, and all authors contributed to revisions.

## Data availability statement

Data, codes and computing environments to reproduce this manuscript will be archived on Zenodo at https://doi.org/xxx and also available on Github at https://github.com/mattocci27/LMAms.

## References

Betancourt, M. 2016. Diagnosing Suboptimal Cotangent Disintegrations in Hamiltonian Monte Carlo. arXiv.

Blonder, B., C. Violle, L. P. Bentley, and B. J. Enquist. 2011. Venation networks and the origin of the leaf economics spectrum. Ecology Letters 14:91–100.

Bonan, G. B., S. Levis, L. Kergoat, and K. W. Oleson. 2002. Landscapes as patches of plant functional types: An integrating concept for climate and ecosystem models. Global Biogeochemical Cycles 16:5-1-5-23.

Carpenter, B., A. Gelman, M. D. Hoffman, D. Lee, B. Goodrich, M. Betancourt, M. Brubaker, J. Guo, P. Li, and A. Riddell. 2017. Stan: A Probabilistic Programming Language. Journal of Statistical Software 76:1–32.

Dantas de Paula, M., M. Forrest, L. Langan, J. Bendix, J. Homeier, A. Velescu, W. Wilcke, and T. Hickler. 2021. Nutrient cycling drives plant community trait assembly and ecosystem functioning in a tropical mountain biodiversity hotspot. New Phytologist 232:551–566.

Dong, N., I. C. Prentice, B. J. Evans, S. Caddy-Retalic, A. J. Lowe, and I. J. Wright. 2017. Leaf nitrogen from first principles: Field evidence for adaptive variation with climate. Biogeosciences 14:481–495.

Dong, N., I. C. Prentice, I. J. Wright, H. Wang, O. K. Atkin, K. J. Bloomfield, T. F. Domingues, S. M. Gleason, V. Maire, Y. Onoda, H. Poorter, and N. G. Smith. 2022. Leaf nitrogen from the perspective of optimal plant function. Journal of Ecology 110:2585–2602.

Evans, J. R., R. Kaldenhoff, B. Genty, and I. Terashima. 2009. Resistances along the CO2 diffusion pathway inside leaves. Journal of Experimental Botany 60:2235–2248.

Falster, D. S., P. B. Reich, D. S. Ellsworth, I. J. Wright, M. Westoby, J. Oleksyn, and T. D. Lee. 2012. Lifetime return on investment increases with leaf lifespan among 10 Australian woodland species. New Phytologist 193:409–419.

Fisher, R. A., C. D. Koven, W. R. L. Anderegg, B. O. Christoffersen, M. C. Dietze, C. E. Farrior, J. A. Holm, G. C. Hurtt, R. G. Knox, P. J. Lawrence, J. W. Lichstein, M. Longo, A. M. Matheny, D. Medvigy, H. C. Muller-Landau, T. L. Powell, S. P. Serbin, H. Sato, J. K. Shuman, B. Smith, A. T. Trugman, T. Viskari, H. Verbeeck, E. Weng, C. Xu, X. Xu, T. Zhang, and P. R. Moorcroft. 2018. Vegetation demographics in Earth System Models: A review of progress and priorities. Global Change Biology 24:35–54.

Fisher, R. A., S. Muszala, M. Verteinstein, P. Lawrence, C. Xu, N. G. McDowell, R. G. Knox, C. Koven, J. Holm, B. M. Rogers, A. Spessa, D. Lawrence, and G. Bonan. 2015. Taking off the training wheels: The properties of a dynamic vegetation model without climate envelopes, CLM4.5(ED). Geoscientific Model Development 8:3593–3619.

Gelman, A., J. B. Carlin, H. S. Stern, D. B. Dunson, A. Vehtari, and D. B. Rubin. 2013. Bayesian Data Analysis, Third Edition. Chapman & Hall/CRC, Boca Raton, FL, USA.

Hidaka, A., and K. Kitayama. 2011. Allocation of foliar phosphorus fractions and leaf traits of tropical tree species in response to decreased soil phosphorus availability on Mount Kinabalu, Borneo. Journal of Ecology 99:849–857.

Huntingford, C., O. K. Atkin, A. Martinez-de la Torre, L. M. Mercado, M. A. Heskel, A. B. Harper, K. J. Bloomfield, O. S. O’Sullivan, P. B. Reich, K. R. Wythers, E. E. Butler, M. Chen, K. L. Griffin, P. Meir, M. G. Tjoelker, M. H. Turnbull, S. Sitch, A. Wiltshire, and Y. Malhi. 2017. Implications of improved representations of plant respiration in a changing climate. Nature Communications 8:1602.

Kikuzawa, K. 1991. A Cost-Benefit Analysis of Leaf Habit and Leaf Longevity of Trees and Their Geographical. The American Naturalist 138:1250–1263.

Kitajima, K., R. A. Cordero, and S. J. Wright. 2013. Leaf life span spectrum of tropical woody seedlings: Effects of light and ontogeny and consequences for survival. Annals of Botany 112:685–699.

Kitajima, K., A. M. Llorens, C. Stefanescu, M. V. Timchenko, P. W. Lucas, and S. J. Wright. 2012. How cellulose-based leaf toughness and lamina density contribute to long leaf lifespans of shade-tolerant species. New Phytologist 195:640–652.

Kitajima, K., S. J. Wright, and J. W. Westbrook. 2016. Leaf cellulose density as the key determinant of inter- and intra-specific variation in leaf fracture toughness in a species-rich tropical forest. Interface Focus 6:20150100.

Landau, W. M. 2021. The targets R package: A dynamic Make-like function-oriented pipeline toolkit for reproducibility and high-performance computing. Journal of Open Source Software 6:2959.

Lemoine, N. P. 2019. Moving beyond noninformative priors: Why and how to choose weakly informative priors in Bayesian analyses. Oikos 128:912–928.

Lichstein, J. W., B. T. Peterson, J. Langebrake, and S. A. McKinley. 2021. Leaf economics of early- and late-successional plants. The American Naturalist.

Lloyd, J., K. Bloomfield, T. F. Domingues, and G. D. Farquhar. 2013. Photosynthetically relevant foliar traits correlating better on a mass vs an area basis: Of ecophysiological relevance or just a case of mathematical imperatives and statistical quicksand? New Phytologist 199:311– 321.

Lusk, C. H., P. B. Reich, R. A. Montgomery, D. D. Ackerly, and J. Cavender-Bares. 2008. Why are evergreen leaves so contrary about shade? Trends in Ecology and Evolution 23:299–303.

Martínez-Vilalta, J., A. Sala, D. Asensio, L. Galiano, G. Hoch, S. Palacio, F. I. Piper, and F. Lloret. 2016. Dynamics of non-structural carbohydrates in terrestrial plants: A global synthesis. Ecological Monographs 86:495–516.

Moorcroft, P. R. 2006. How close are we to a predictive science of the biosphere? Trends in Ecology & Evolution 21:400–407.

Moorcroft, P. R., G. C. Hurtt, and S. W. Pacala. 2001. A Method for Scaling Vegetation Dynamics: The Ecosystem Demography Model (ED). Ecological Monographs 71:557–585.

Onoda, Y., F. Schieving, and N. P. R. Anten. 2008. Effects of light and nutrient availability on leaf mechanical properties of *Plantago Major*: A conceptual approach. Annals of Botany 101:727–736.

Onoda, Y., F. Schieving, and N. P. R. Anten. 2015. A novel method of measuring leaf epidermis and mesophyll stiffness shows the ubiquitous nature of the sandwich structure of leaf laminas in broad-leaved angiosperm species. Journal of Experimental Botany 66:2487–2499.

Onoda, Y., M. Westoby, P. B. Adler, A. M. F. Choong, F. J. Clissold, J. H. C. Cornelissen, S. Díaz, N. J. Dominy, A. Elgart, L. Enrico, P. V. A. Fine, J. J. Howard, A. Jalili, K. Kitajima, H. Kurokawa, C. McArthur, P. W. Lucas, L. Markesteijn, N. Pérez-Harguindeguy, L. Poorter, L. Richards, L. S. Santiago, E. E. Sosinski, S. A. Van Bael, D. I. Warton, I. J. Wright, S. Joseph Wright, and N. Yamashita. 2011. Global patterns of leaf mechanical properties. Ecology Letters 14:301– 312.

Onoda, Y., I. J. Wright, J. R. Evans, K. Hikosaka, K. Kitajima, Ü. Niinemets, H. Poorter, T. Tosens, and M. Westoby. 2017. Physiological and structural tradeoffs underlying the leaf economics spectrum. New Phytologist 214:1447–1463.

Osada, N., H. Takeda, A. Furukawa, and M. Awang. 2001. Leaf dynamics and maintenance of tree crowns in a malaysian rain forest stand. Journal of Ecology 89:774–782.

Osnas, J. L. D., M. Katabuchi, K. Kitajima, S. J. Wright, P. B. Reich, S. A. Van Bael, N. J. B. Kraft, M. J. Samaniego, S. W. Pacala, and J. W. Lichstein. 2018. Divergent drivers of leaf trait variation within species, among species, and among functional groups. Proceedings of the National Academy of Sciences of the United States of America 115:5480–5485.

Osnas, J. L. D., J. W. Lichstein, P. B. Reich, and S. W. Pacala. 2013. Global leaf trait relationships: Mass, area, and the leaf economics spectrum. Science 340:741–744.

Poorter, H., Ü. Niinemets, L. Poorter, I. J. Wright, and R. Villar. 2009. Causes and consequences of variation in leaf mass per area (LMA): A meta-analysis. New Phytologist 182:565–588.

R Core Team. 2022. R: A language and environment for statistical computing. manual, R Foundation for Statistical Computing, Vienna, Austria.

Reich, P. B. 2014. The world-wide ’fast-slow’ plant economics spectrum: A traits manifesto. Journal of Ecology 102:275–301.

Russo, S. E., and K. Kitajima. 2016. The Ecophysiology of Leaf Lifespan in Tropical Forests: Adaptive and Plastic Responses to Environmental Heterogeneity. Pages 357–383 in G. Goldstein and S. L. Santiago, editors. Tropical Tree Physiology. Springer International Publishing, Cham.

Sakschewski, B., W. von Bloh, A. Boit, L. Poorter, M. Peña-Claros, J. Heinke, J. Joshi, and K. Thonicke. 2016. Resilience of Amazon forests emerges from plant trait diversity. Nature Climate Change 6:1032–1036.

Sakschewski, B., W. von Bloh, A. Boit, A. Rammig, J. Kattge, L. Poorter, J. Peñuelas, and K. Thonicke. 2015. Leaf and stem economics spectra drive diversity of functional plant traits in a dynamic global vegetation model. Global Change Biology 21:2711–2725.

Scheiter, S., L. Langan, and S. I. Higgins. 2013. Next-generation dynamic global vegetation models: Learning from community ecology. New Phytologist 198:957–969.

Shipley, B., M. J. Lechowicz, I. Wright, and P. B. Reich. 2006. Fundamental trade-offs generating the worldwide leaf economics spectrum. Ecology 87:535–541.

Tcherkez, G., P. Gauthier, T. N. Buckley, F. A. Busch, M. M. Barbour, D. Bruhn, M. A. Heskel, X. Y. Gong, K. Y. Crous, K. Griffin, D. Way, M. Turnbull, M. A. Adams, O. K. Atkin, G. D. Farquhar, and G. Cornic. 2017. Leaf day respiration: Low CO2 flux but high significance for metabolism and carbon balance. New Phytologist 216:986–1001.

Terashima, I., Y. T. Hanba, D. Tholen, and U. Niinemets. 2011. Leaf Functional Anatomy in Relation to Photosynthesis. Plant Physiology 155:108–116.

Vehtari, A., A. Gelman, and J. Gabry. 2017. Practical Bayesian model evaluation using leave-one-out cross-validation and WAIC. Statistics and Computing:1413–1432.

Verheijen, L. M., V. Brovkin, R. Aerts, G. Bönisch, J. H. C. Cornelissen, J. Kattge, P. B. Reich, I. J. Wright, and P. M. van Bodegom. 2013. Impacts of trait variation through observed trait– climate relationships on performance of an Earth system model: A conceptual analysis. Biogeosciences 10:5497–5515.

Villar, R., and J. Merino. 2001. Comparison of leaf construction costs in woody species with differing leaf life-spans in contrasting ecosystems. New Phytologist 151:213–226.

Westoby, M., P. B. Reich, and I. J. Wright. 2013. Understanding ecological variation across species: Area-based vs mass-based expression of leaf traits. New Phytologist 199:322– 323.

Westoby, M., D. Warton, and P. B. Reich. 2000. The Time Value of Leaf Area. The American Naturalist 155:649–656.

Williams, K., C. B. Field, and H. A. Mooney. 1989. Relationships Among Leaf Construction Cost, Leaf Longevity, and Light Environment in Rain-Forest Plants of the Genus Piper. The American Naturalist 133:198–211.

Wright, I. J., P. B. Reich, M. Westoby, D. D. Ackerly, Z. Baruch, F. Bongers, J. Cavender-Bares, T. Chapin, J. H. C. Cornellssen, M. Diemer, J. Flexas, E. Garnier, P. K. Groom, J. Gulias, K. Hikosaka, B. B. Lamont, T. Lee, W. Lee, C. Lusk, J. J. Midgley, M. L. Navas, Ü. Niinemets, J. Oleksyn, H. Osada, H. Poorter, P. Pool, L. Prior, V. I. Pyankov, C. Roumet, S. C. Thomas, M. G. Tjoelker, E. J. Veneklaas, and R. Villar. 2004. The worldwide leaf economics spectrum. Nature 428:821–827.

Wright, S. J., V. Horlyck, Y. Basset, H. Barrios, A. Bethancourt, S. Bohlman, G. Gilbert, G. Goldstein, E. A. Graham, K. Kitajima, M. T. Lerdau, F. C. Meinzer, F. Ødegaard, D. R. Reynolds, D. W. Roubik, S. Sakai, M. Samaniego, J. P. Sparks, S. Van Bael, K. Winter, and G. Zotz. 2003. Tropical Canopy Biology Program, Republic of Panama. Pages 137–155 *in* Y. Basset, V. Horlyck, and S. J. Wright, editors. Studying Forest Canopies from Above: The International Canopy Crane Network. Panama.

Wullschleger, S. D., H. E. Epstein, E. O. Box, E. S. Euskirchen, S. Goswami, C. M. Iversen, J. Kattge, R. J. Norby, P. M. Van Bodegom, and X. Xu. 2014. Plant functional types in Earth system models: Past experiences and future directions for application of dynamic vegetation models in high-latitude ecosystems. Annals of Botany 114:1–16.

Würth, M. K. R., S. Peláez-Riedl, S. J. Wright, and C. Körner. 2005. Non-structural carbohydrate pools in a tropical forest. Oecologia 143:11–24.

Xu, X., D. Medvigy, S. Joseph Wright, K. Kitajima, J. Wu, L. P. Albert, G. A. Martins, S. R. Saleska, and S. W. Pacala. 2017. Variations of leaf longevity in tropical moist forests predicted by a trait-driven carbon optimality model. Ecology Letters 20:1097–1106.

