## Appendix for "Decomposing leaf mass into metabolic and structural components explains divergent patterns of trait variation within and among plant species"

**References** **21**

### Appendix S1: Prior information

The logarithms of $A_{\text{area}}$, $R_{\text{area}}$, and LL for leaf sample $i$ were assumed to have a multivariate normal distribution:

$$\begin{pmatrix} \ln(A_{\text{area}\,i}) \\ \ln(R_{\text{area}\,i}) \\ \ln(\text{LL}_i) \end{pmatrix} \sim \text{MVN}\left( \begin{pmatrix} \text{E}[A_{\text{area}\,i}] \\ \text{E}[R_{\text{area}\,i}] \\ \text{E}[\text{LL}_i] \end{pmatrix}, \Sigma \right) \tag{S1}$$

where E[$\cdot$] indicates expected value and $\Sigma$ indicates a covariance matrix. Expected values are based on Eqs. 2- 4 in the main text.

We used non-informative or weakly informative prior distributions (Lemoine, 2019). The covariance matrix in Eq. S1 was decomposed as $\Sigma = \text{diag}(\sigma)\Omega\text{diag}(\sigma) = \text{diag}(\sigma)LL'\text{diag}(\sigma)$ using a Cholesky decomposition, where $\sigma$ is a vector of $\sigma_1$, $\sigma_2$, and $\sigma_3$; $\Omega$ is a correlation matrix of $\rho_{12}$, $\rho_{13}$, and $\rho_{23}$; and **L** is a lower triangular matrix. Instead of assigning prior distributions on $\Sigma$ directly, priors were assigned on $\sigma$ and **L** to avoid a strong dependence between $\sigma$ and $\Omega$ (Alvarez et al., 2014; Lewandowski et al., 2009). A prior for **L** was specified as a so-called LKJ distribution with shape parameter 2 (Lewandowski et al., 2009), which is weakly informative for the correlation matrix. A prior for $\sigma$ was specified as a Half-Cauchy distribution with location 0 and scale 2.5, which is weakly informative and allows for occasional large coefficients while still performing a reasonable amount of shrinkage for coefficients near zero (Gelman et al., 2008). Priors for $\alpha_{0,m,s}$, $\beta_{0,m,s}$, and $\gamma_{0,m,s}$ in Eqs. 2- 4 were weakly informative and specified as normal distributions with mean 0 and standard deviation 5. Priors for $f_i$ in Eqs. 1- 4 were non-informative and specified as uniform distributions with range (0, 1).

### Appendix S2: Model tests with randomized data (including Table AS1 and Figure AS1)

We generated 10 randomized datasets for each of the GLOPNET and the Panama datasets by randomly shuffling each trait value across leaf samples, and we fit the best models (Table 1) to the randomized data.

Most of the model results obtained from the randomized datasets did not convergent based on the Gelman-Rubin statistic or showed divergent transitions (Table AS1), indicating that models fit to randomized data do not provide reliable inferences (Betancourt, 2016).

#### Table AS1

- No_large_Rhat: The number of parameters (including transformed parameters) that shows Rhat (Gelman-Rubin statistic) greater than 1.1.
- No_divergence: The number of iterations that shows divergent transitions.

| Data | Simulation_ID | No_large_Rhat | No_divergence |
|---|---|---:|---:|
| GLOPNET | sim-01 | 4 | 13 |
| GLOPNET | sim-02 | 4 | 0 |
| GLOPNET | sim-03 | 0 | 0 |
| GLOPNET | sim-04 | 0 | 0 |
| GLOPNET | sim-05 | 0 | 1 |
| GLOPNET | sim-06 | 0 | 1 |
| GLOPNET | sim-07 | 12 | 0 |
| GLOPNET | sim-08 | 4 | 0 |
| GLOPNET | sim-09 | 0 | 0 |
| GLOPNET | sim-10 | 104 | 1 |
| Panama | sim-01 | 4 | 131 |
| Panama | sim-02 | 64 | 23 |
| Panama | sim-03 | 397 | 28 |
| Panama | sim-04 | 0 | 15 |
| Panama | sim-05 | 7 | 58 |
| Panama | sim-06 | 79 | 4 |
| Panama | sim-07 | 7 | 122 |
| Panama | sim-08 | 160 | 131 |
| Panama | sim-09 | 451 | 579 |
| Panama | sim-10 | 601 | 148 |

### Figure AS1

Although many parameters in models fit to randomized data had large Rhat values (Table AS1), which suggests that the posterior distributions did not converge, we nevertheless examined the regression coefficients to evaluate if our model framework is prone to overfitting. Figure AS1 shows the regression coefficients for the randomized GLOPNET data ($\alpha_{0,m,s}$, $\beta_{0,s}$, and $\gamma_{0,m,s}$ in Eqs. 2- 4 in the main text). There are 10 independent simulations (randomizations) in total. Points and lines indicate posterior medians and 95% credible intervals (CIs), respectively. Although intercepts were significant, the scaling parameters ($\alpha_{m,s}$, $\beta_s$, and $\gamma_{m,s}$) obtained for the randomized datasets were not significantly different from zero. These tests with randomized data indicate that our model is not inherently prone to overfitting or to producing patterns from noise.

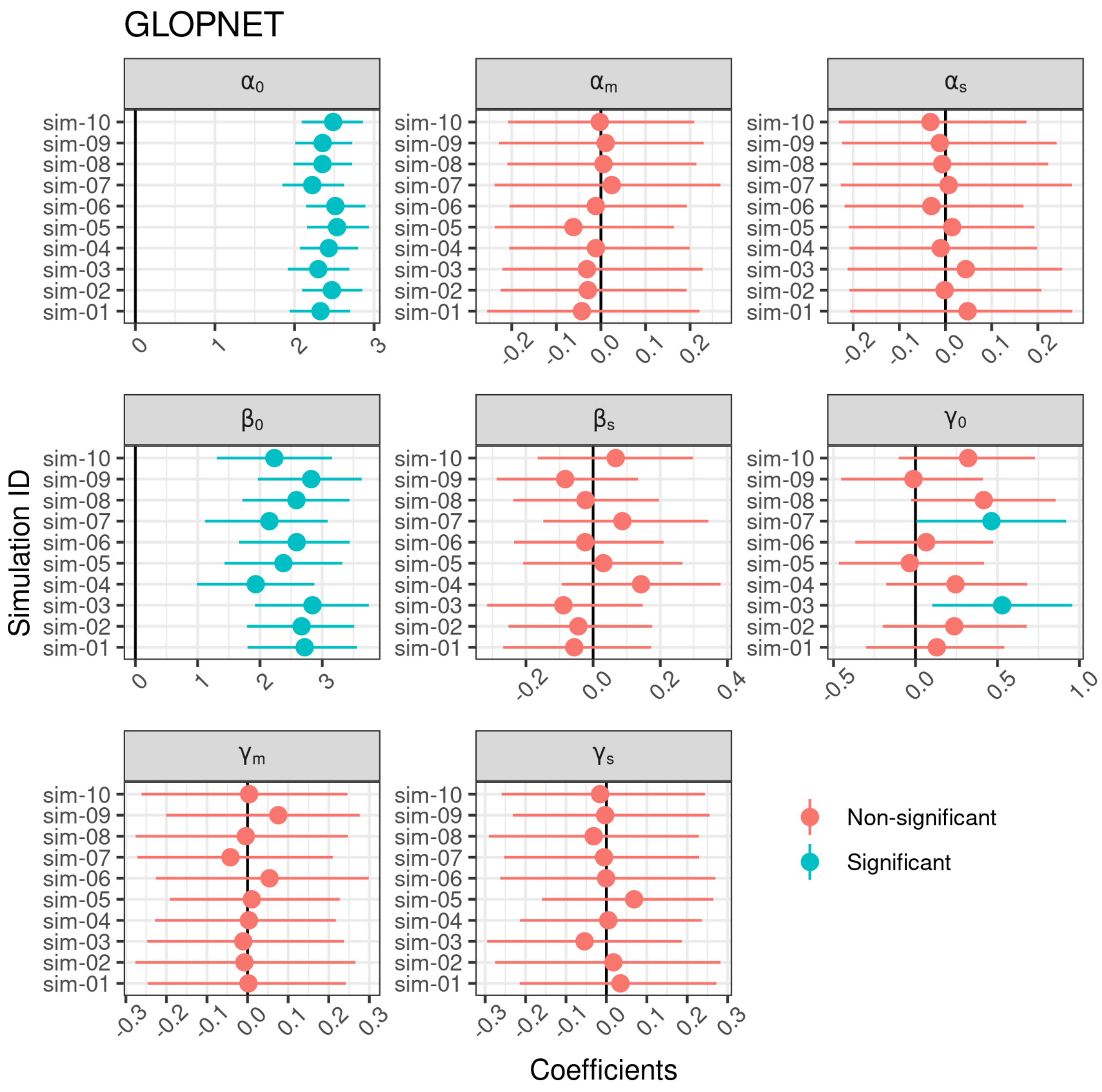

#### Figure AS2

Figure AS2 showsWe also checked the regression coefficients for the randomized Panama dataset ($\alpha_{0,m}$, $\beta_{0,s}$, $\gamma_{0,m,s}$, and $\theta$ in Eqs. 2- 4 in the main text). Details as for Figure AS1. The scaling parameters ($\alpha_m$, $\beta_s$, and $\gamma_{m,s}$) and the effect of light on leaf lifespan ($\theta$) were not significantly different from zero in the randomized datasets did not show any patterns. Again, these tests with randomized data indicate that our model is not inherently prone to overfitting or to producing patterns from noise.

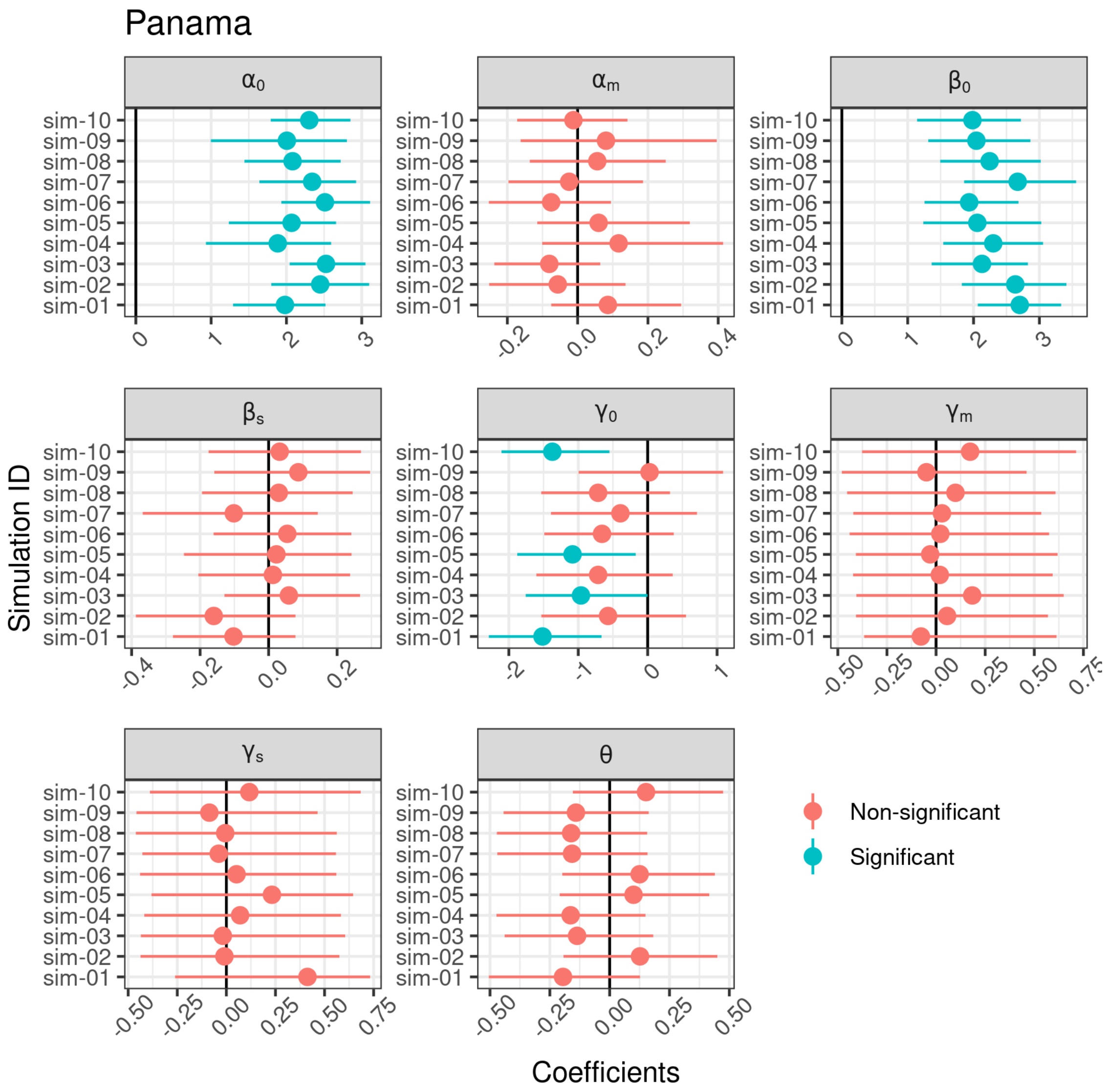

### Appendix S3: Simulated datasets to study relationships between photosynthetic capacity and LMA

To create simulated datasets to study relationships between photosynthetic capacity and LMA , we generated log-normally distributed LMAm and LMAs values by sampling ln(LMAm) and ln(LMAs) from normal distributions.

For GLOPNET and Panama sun leaves, results from the LMAm-LMAs models (Eqs. 2- 4) indicated little correlation between LMAm and LMAs, so we used univariate normal distributions (N) to generate the LMAm and LMAs samples:

$$\ln(\mathrm{LMAm}) \sim \mathrm{N}(\ln(\mu_m), \sigma_m) \qquad \text{(S3.1)}$$

$$\ln(\mathrm{LMAs}) \sim \mathrm{N}(\ln(\mu_s), \sigma_s) \qquad \text{(S3.2)}$$

where $\ln(\mu_m)$ and $\ln(\mu_s)$ are the means of LMAm and LMAs, respectively, on the log-scale and $\sigma_m$ and $\sigma_s$ are the corresponding standard deviations. The parameters $\ln(\mu_m)$, $\ln(\mu_s)$, $\sigma_m$ and $\sigma_s$ were all estimated from the posterior medians of LMAm and LMAs.

For Panama shade leaves, LMAm-LMAs model results indicated a negative correlation between LMAm and LMAs, so we used a multivariate normal distribution (MVN) to generate the LMAm and LMAs samples:

$$\begin{bmatrix}\ln(\mathrm{LMAm}) \\ \ln(\mathrm{LMAs})\end{bmatrix} \sim \mathrm{MVN}\begin{bmatrix}\ln(\mu_m) & \\ \ln(\mu_s) & , \Sigma\end{bmatrix} \qquad \text{(S3.3)}$$

$$\Sigma = \begin{bmatrix}\sigma_m^2 & \rho\sigma_m\sigma_s \\ \rho\sigma_m\sigma_s & \sigma_s^2\end{bmatrix} \qquad \text{(S3.4)}$$

where $\Sigma$ is the covariance matrix of ln(LMAm) and ln(LMAs), and $\rho$ is the correlation coefficient between ln(LMAm) and ln(LMAs). The parameters in $\Sigma$ were all estimated from the posterior medians of LMAm and LMAs.

To create simulated datasets (each with a sample size of 100) in which LMAs accounted for different fractions of the total LMA variance, we used the predictions (posterior medians) of LMAm and LMAs for each leaf sample from the best models (Table 2) to estimate $\mu_m$, $\mu_s$, $\sigma_m$, and $\rho$; and we varied $\sigma_s$ from ln(1.01) to ln(10) across simulated datasets. For each of the 100 leaves in a given simulated dataset, $A_{\mathrm{area}}$ was calculated according to Eq. 2.

Parameter values were $\mu_m$ = 56.5, $\mu_s$ = 61.2, $\sigma_m$ = 0.83, $\alpha_0$ = 1.77, $\alpha_m$ = 0.28, and $\alpha_s$ = -0.13 for GLOPNET; $\mu_m$ = 47.3, $\mu_s$ = 30.1, $\sigma_m$ = 1.58, $\alpha_0$ = 0.34, $\alpha_m$ = 0.56, and $\alpha_s$ = 0 for Panama sun leaves; and $\mu_m$ = 7.6, $\mu_s$ = 26.7, $\sigma_m$ = 1.95, $\alpha_0$ = 0.34, $\alpha_m$ = 0.56, $\alpha_s$ = 0, and $\rho$ = -0.47 for Panama shade leaves.

For each simulated dataset, we quantified the mass-dependence ($b$) using the ordinary least squares regression (log-log) form of Eq. 5. We repeated these steps 1000 times for each set of parameter values.

We also used Eqs. S3.1-S3.2 to create the simulated LMA datasets in Fig. 1, which is based on our analysis of GLOPNET data. $A_{\text{area}}$ values were generated using Eq. 2 and Eq. S1. Based on our GLOPNET results, the standard deviations of ln(LMAs) and ln($A_{\text{area}}$) were set to 1 and 0.31, respectively, and other GLOPNET parameter values are listed above.

### Figures

#### Fig. S1

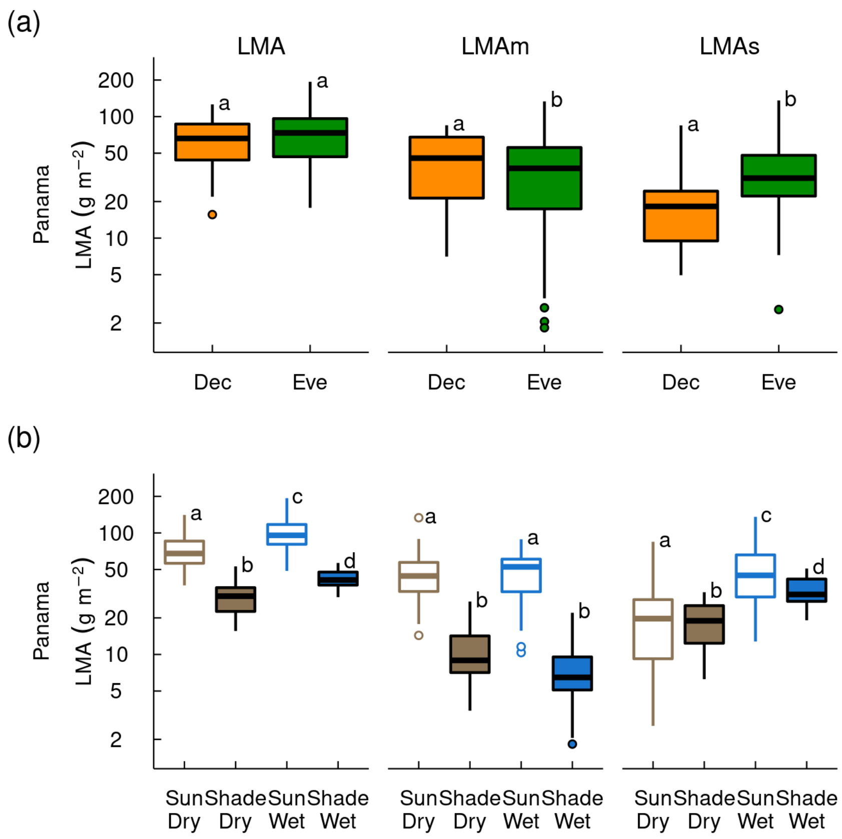

**Fig. S1**: Boxplots comparing leaf mass per area (LMA), photosynthetic leaf mass per area (LMAm; posterior medians), and structural leaf mass per area (LMAs; posterior medians) across (a) deciduous (Dev) and evergreen (Eve) leaves and (b) sites (wet and dry) and canopy strata (sun and shade) in Panama. The results shown here include all leaves in the Panama dataset, whereas Figs. 6- 7 in the main text only include Panama species for which both sun and shade leaves were available. Boxplot symbols as in Figs. 6- 7. Groups sharing the same letters are not significantly different ($P > 0.05$; t-tests). Estiamtes are from the best Panama model (Table 1).

#### Fig. S2

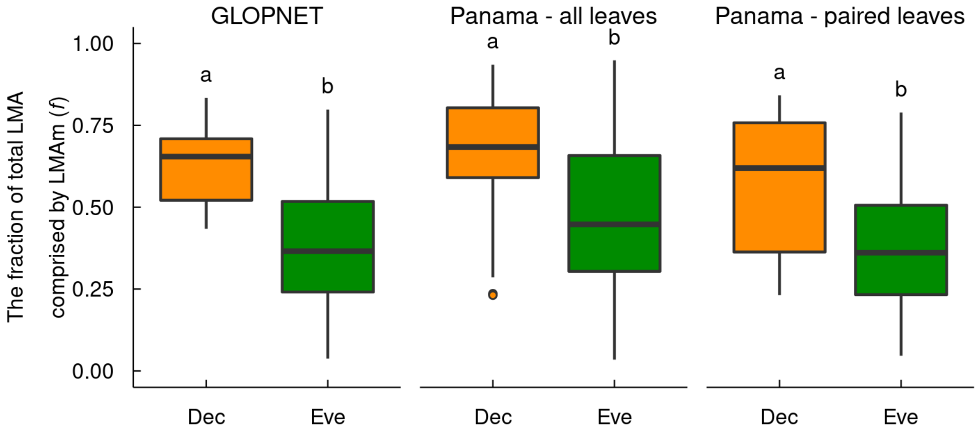

**Fig. S2**: Boxplots comparing posterior medians of the latent variable $f$ (the fraction of total LMA comprised by LMAm) across deciduous (Dev) and evergreen (Eve) leaves. Left: GLOPNET dataset. Middle: All leaf samples in the Panama dataset. Right: Paired leaf samples in the Panama dataset (species for which both sun and shade leaves were available). Note that LMAm $= f \times$ LMA, and LMAs $= (1 - f) \times$ LMA. Boxplot symbols as in Fig. 6. Groups sharing the same letters are not significantly different ($P > 0.05$; t-tests). Estiamtes are from the best GLOPNET and Panama models (Table 1).

### Fig. S3

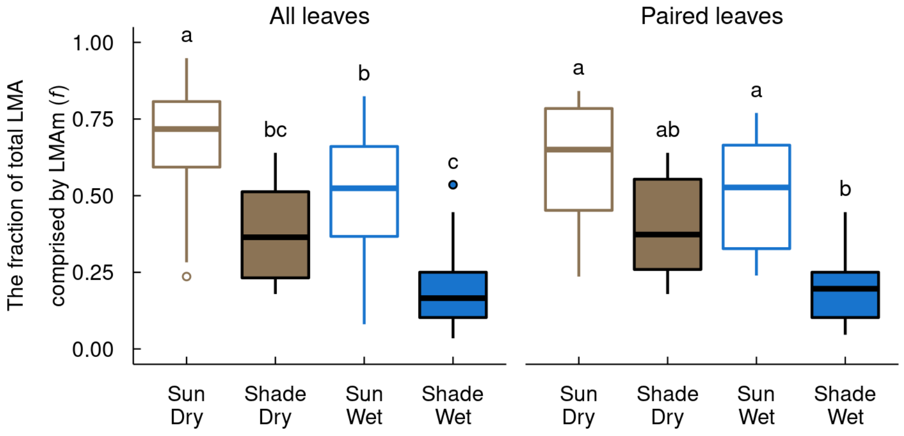

**Fig. S3**: Boxplots comparing posterior medians of the latent variable $f$ (the fraction of total LMA comprised by LMAm) across sites (wet and dry) and canopy strata (sun and shade) in Panama. Left: All leaf samples in the Panama dataset. Right: Paired leaf samples in the Panama dataset (species for which both sun and shade leaves were available). Note that LMAm = $f \times$ LMA, and LMAs = $(1 - f) \times$ LMA. Boxplot symbols as in Fig. 7. Groups sharing the same letters are not significantly different ($P > 0.05$; t-tests). Estiamtes are from the best GLOPNET and Panama models (Table 1).

### Fig. S4

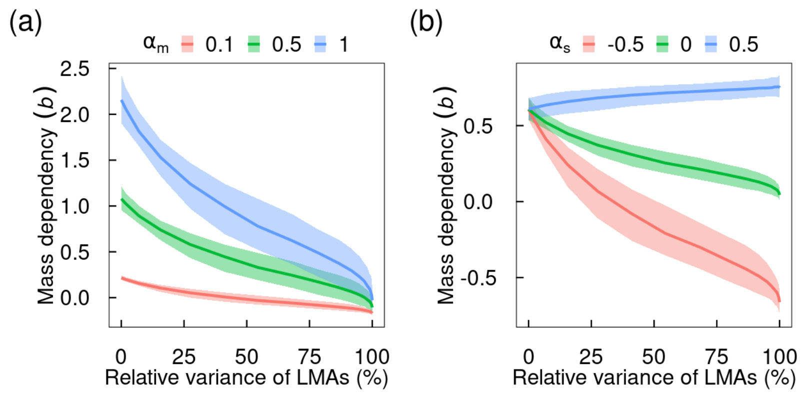

**Fig. S4**: The relationships between mass dependency of $A_{max}$ (*b* in Eq. 5 in the main text) and LMAs variance (relative to total LMA variance; Eq. 6) for the different values of the scaling exponents $\alpha_m$ and $\alpha_s$ (Eq. 2). (a) The scaling exponent $\alpha_m$ varies from 0.1 to 1.0 while the scaling exponent $\alpha_s$ is constant ($\alpha_s$ = -0.13). (b) The scaling exponent $\alpha_s$ vary from -0.5 to 0.5 while the scaling exponent $\alpha_m$ is constant ($\alpha_m$ = 0.28). Solid lines indicate simulated medians and shaded regions indicate 95% confidence intervals. Empirical estimates of *b* are typically between 0 and 1 (Osnas et al., 2018). $A_{max}$ is primarily mass-dependent if *b* > 0.5, and primarily area-dependent if 0.5 > *b* > 0 (Osnas et al., 2018). Parameter values are based on the best GLOPNET model (Table 1) and Appendix S3: $\alpha_0$ = 1.77, $\mu_m$ = 56.5, $\mu_s$ = 61.2, and $\sigma_m$ = 0.83.

### Fig. S5

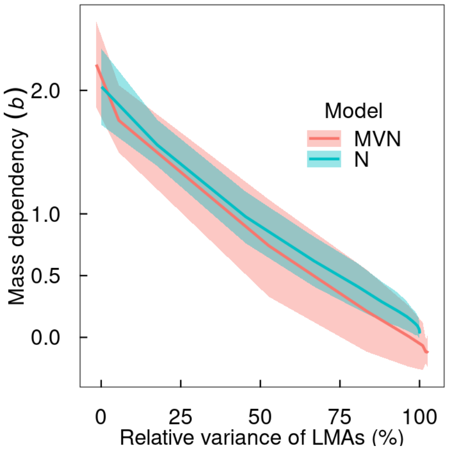

**Fig. S5**: The relationships between mass dependency of $A_{max}$ (*b* in Eq. 5 in the main text) and LMAs variance (relative to total LMA variance; Eq. 6) for simulated datasets generated from a normal distribution (N) vs. a multivariate normal distribution (MVN). Parameter values are based on the best Panama model (Table 1) and Appendix S3: $\alpha_0$ = 0.34, $\alpha_m$ = 0.56, $\alpha_s$ = 0, $\mu_m$ = 7.6, $\mu_s$ = 26.7, $\sigma_m$ = 1.95, and $\rho$ = -0.47.

### Fig. S6

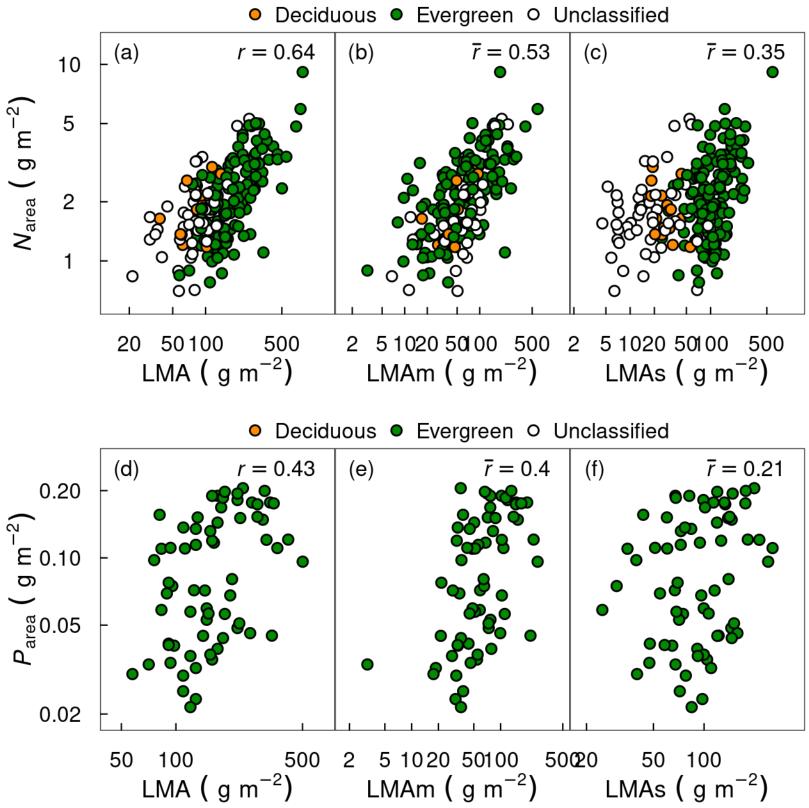

**Fig. S6**: Measured traits related to photosynthesis and metabolism (nitrogen and phosphorus per-unit leaf area; $N_{\text{area}}$ and $P_{\text{area}}$) are positively correlated with LMA and with estimates (posterior medians) of the metabolic and structural LMA components (LMAm and LMAs, respectively) in the GLOPNET dataset. LMAm yields more consistent relationships compared to LMA and LMAs; e.g., evergreen and deciduous leaves align along a single relationship in panel b, but not in panels a or c. Pearson correlation coefficients ($r$) for LMA (left column) and posterior medians of Pearson correlation coefficients ($\bar{r}$) for LMAm (middle column) and LMAs (right column) are shown.

### Fig. S7

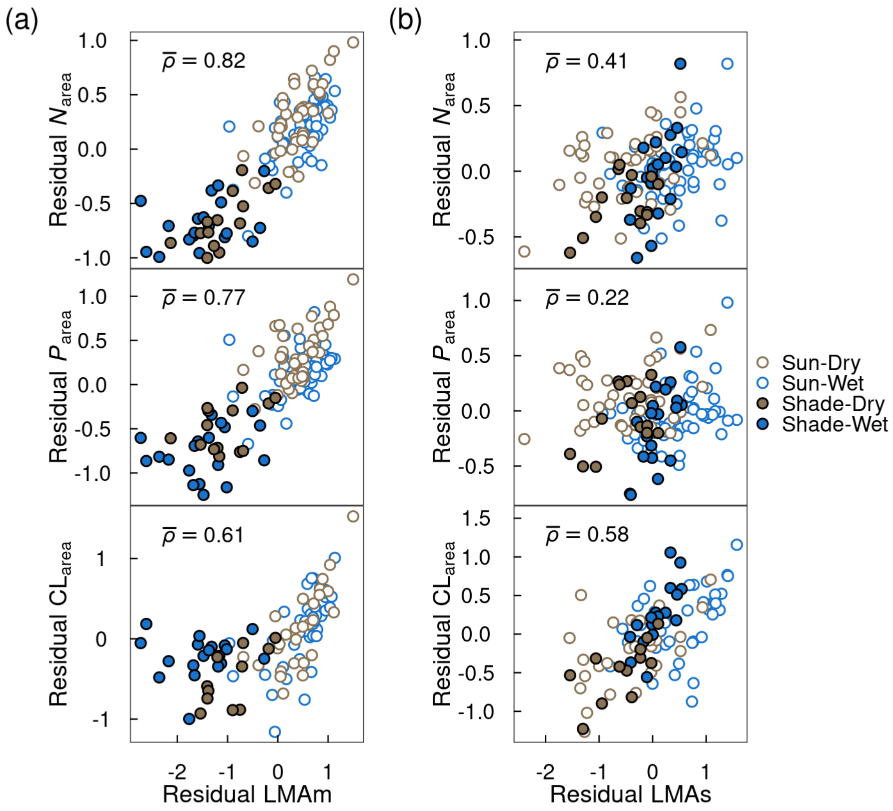

**Fig. S7**: Partial regression plots for nitrogen, phosphorus and cellulose per-unit leaf area ($N_{area}$, $P_{area}$ and $CL_{area}$). (a) LMAs variation is controlled. (b) LMAm variation is controlled. The partial regression plots show separation between sun and shade when controlling for LMAs variation (i.e., LMAs does not explain the sun/shade difference), but overlapping distributions of sun and shade when controlling for LMAm variation (i.e., LMAm does explain the sun/shade difference). Posterior medians of partial correlation coefficients ($\bar{\rho}$) are shown.

### Fig. S8

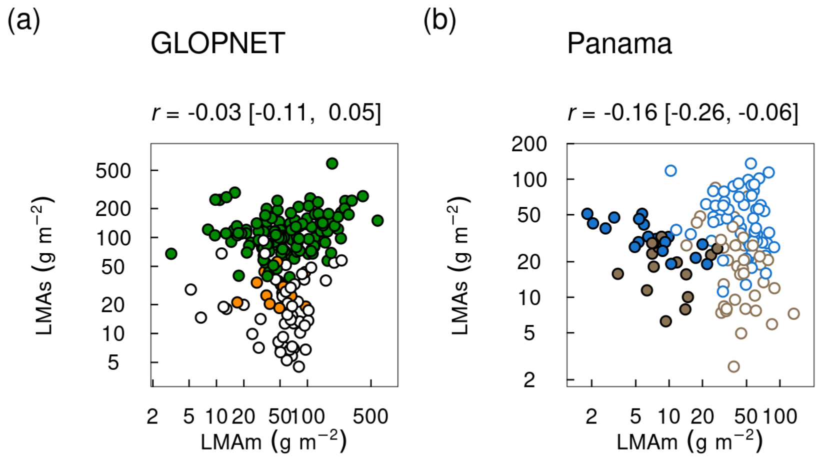

**Fig. S8** Pearson correlation coefficients for posterior medians of LMAm vs LMAs in the (a) GLOPNET and (b) Panama datasets. The non-significant or weak *r* values indicate that a single axis could not accurately represent the two-dimensional space. Symbols as in main text Figs. 2- 3.

### Appendix S4: Stan code

#### Stan code for the GLOPNET dataset

The best model for the GLOPNET dataset is LMAm-LMAs with the constraint of $\beta_m$ = 0.

```
//NOTE: THIS STAN CODE IS GENERATED VIA "update.py"
data {
  int<lower=0> N;
  vector<lower=0>[N] LMA;
  vector<lower=0>[N] A;
  vector<lower=0>[N] R;
  vector<lower=0>[N] LL;
}

transformed data {
  vector[N] log_A;
  vector[N] log_LL;
  vector[N] log_R;
  matrix[N, 3] obs;
  vector[N] intercept;
  for (n in 1:N)
    intercept[n] = 1;
  log_A = log(A);
  log_LL = log(LL);
  log_R = log(R);
  // use net photosynthesis (A) instead of gross (A + R)
  obs = append_col(append_col(log_A, log_LL), log_R);
}

parameters {
  real a0;
  real am;
  real as;
  real b0;
  real bs;
  real g0;
  real gm;
  real gs;
  vector<lower=0, upper=1>[N] p;
  vector<lower=0>[3] L_sigma;
  cholesky_factor_corr[3] L_Omega;
}
```

```
transformed parameters {
  matrix[N, 3] Mu;
  matrix[3, 3] Z;
  matrix[N, 3] X;
  Z[1, 1] = a0;
  Z[1, 2] = b0;
  Z[1, 3] = g0;
  Z[2, 1] = am;
  Z[2, 2] = 0;
  Z[2, 3] = gm;
  Z[3, 1] = as;
  Z[3, 2] = bs;
  Z[3, 3] = gs;

  //log_LMAm = log(LMA) + log(p);
  //log_LMAs = log(LMA) + log(1 - p);
  //X = append_col(append_col(append_col(intercept,
   ↪  log_LMAm), log_LMAs), leaf);
  X = append_col(append_col(intercept, log(LMA) + log(p)),
 ↪  log(LMA) + log(1 - p));
  Mu = X * Z;
}

model {
  // priors
  a0 ~ normal(0, 5);
  b0 ~ normal(0, 5);
  g0 ~ normal(0, 5);
  am ~ normal(0, 5);
  bs ~ normal(0, 5);
  gm ~ normal(0, 5);
  gs ~ normal(0, 5);
  as ~ normal(0, 5);
  p ~ beta(1, 1);
  L_Omega ~ lkj_corr_cholesky(2);
  L_sigma ~ cauchy(0, 2.5);

  // model
  for (i in 1:N)
     target += multi_normal_cholesky_lpdf(obs[i,] | Mu[i,],
      ↪  diag_pre_multiply(L_sigma, L_Omega));
}
```

```
generated quantities {
  vector[N] log_lik;
  real<lower=-1, upper=1> rho12;
  real<lower=-1, upper=1> rho23;
  real<lower=-1, upper=1> rho13;
  cov_matrix[3] Sigma;
  vector[N] log_LMAm;
  vector[N] log_LMAs;
  log_LMAm = log(LMA) + log(p);
  log_LMAs = log(LMA) + log(1 - p);
  Sigma = diag_pre_multiply(L_sigma, L_Omega)
     * diag_post_multiply(L_Omega', L_sigma);
  rho12 = Sigma[1, 2] * inv(L_sigma[1] * L_sigma[2]);
  rho23 = Sigma[2, 3] * inv(L_sigma[2] * L_sigma[3]);
  rho13 = Sigma[1, 3] * inv(L_sigma[1] * L_sigma[3]);
  for (i in 1:N)
   log_lik[i] = multi_normal_cholesky_lpdf(obs[i,] | Mu[i,],
 ↪  diag_pre_multiply(L_sigma, L_Omega));
 }
```

#### Stan code for the Panama dataset

The best model for the Panama dataset is LMAm-LMAs-light with the constraint of $\alpha_s = 0$ and $\beta_m = 0$.

```
//NOTE: THIS STAN CODE IS GENERATED VIA "update.py"
data {
  int<lower=0> N;
  vector<lower=0>[N] LMA;
  vector<lower=0>[N] A;
  vector<lower=0>[N] R;
  vector<lower=0>[N] LL;
  vector<lower=0>[N] leaf;
}

transformed data {
  vector[N] log_A;
  vector[N] log_LL;
  vector[N] log_R;
  matrix[N, 3] obs;
```

```
  vector[N] intercept;
  for (n in 1:N)
    intercept[n] = 1;
  log_A = log(A);
  log_LL = log(LL);
  log_R = log(R);
  // use net photosynthesis (A) instead of gross (A + R)
  obs = append_col(append_col(log_A, log_LL), log_R);
}

parameters {
  real a0;
  real am;
  real b0;
  real bs;
  real g0;
  real gm;
  real gs;
  real theta;
  vector<lower=0, upper=1>[N] p;
  vector<lower=0>[3] L_sigma;
  cholesky_factor_corr[3] L_Omega;
}

transformed parameters {
  matrix[N, 3] Mu;
  matrix[4, 3] Z;
  matrix[N, 4] X;
  Z[1, 1] = a0;
  Z[1, 2] = b0;
  Z[1, 3] = g0;
  Z[2, 1] = am;
  Z[2, 2] = 0;
  Z[2, 3] = gm;
  Z[3, 1] = 0;
  Z[3, 2] = bs;
  Z[3, 3] = gs;
  Z[4, 1] = 0;
  Z[4, 2] = theta;
  Z[4, 3] = 0;

  //log_LMAm = log(LMA) + log(p);
  //log_LMAs = log(LMA) + log(1 - p);
```

```
  //X = append_col(append_col(append_col(intercept,
   ↪  log_LMAm), log_LMAs), leaf);
  X = append_col(append_col(append_col(intercept,
    log(LMA) + log(p)),
    log(LMA) + log(1 - p)),
    leaf);
  Mu = X * Z;
}

model {
  // priors
  a0 ~ normal(0, 5);
  b0 ~ normal(0, 5);
  g0 ~ normal(0, 5);
  am ~ normal(0, 5);
  bs ~ normal(0, 5);
  gm ~ normal(0, 5);
  gs ~ normal(0, 5);
  theta ~ normal(0, 5);
  p ~ beta(1, 1);
  L_Omega ~ lkj_corr_cholesky(2);
  L_sigma ~ cauchy(0, 2.5);

  // model
  for (i in 1:N)
     target += multi_normal_cholesky_lpdf(obs[i,] | Mu[i,],
      ↪  diag_pre_multiply(L_sigma, L_Omega));
}

generated quantities {
  vector[N] log_lik;
  real<lower=-1, upper=1> rho12;
  real<lower=-1, upper=1> rho23;
  real<lower=-1, upper=1> rho13;
  cov_matrix[3] Sigma;
  vector[N] log_LMAm;
  vector[N] log_LMAs;
  log_LMAm = log(LMA) + log(p);
  log_LMAs = log(LMA) + log(1 - p);
  Sigma = diag_pre_multiply(L_sigma, L_Omega)
     * diag_post_multiply(L_Omega', L_sigma);
  rho12 = Sigma[1, 2] * inv(L_sigma[1] * L_sigma[2]);
  rho23 = Sigma[2, 3] * inv(L_sigma[2] * L_sigma[3]);
```

```
  rho13 = Sigma[1, 3] * inv(L_sigma[1] * L_sigma[3]);
  for (i in 1:N)
   log_lik[i] = multi_normal_cholesky_lpdf(obs[i,] | Mu[i,],
↪  diag_pre_multiply(L_sigma, L_Omega));
}
```

### References

Alvarez, I., Niemi, J., & Simpson, M. (2014). *Bayesian inference for a covariance matrix* (pp. 1–12). https://doi.org/10.1214/aos/1176348885

Betancourt, M. (2016). Diagnosing Suboptimal Cotangent Disintegrations in Hamiltonian Monte Carlo. *arXiv*. https://doi.org/10.48550/arXiv.1604.00695

Gelman, A., Jakulin, A., Pittau, M. G., & Su, Y. S. (2008). A weakly informative default prior distribution for logistic and other regression models. *Annals of Applied Statistics*, *2*(4), 1360–1383. https://doi.org/10.1214/08-AOAS191

Lemoine, N. P. (2019). Moving beyond noninformative priors: Why and how to choose weakly informative priors in Bayesian analyses. *Oikos*, *128*(7), 912–928. https://doi.org/10.1111/oik.05985

Lewandowski, D., Kurowicka, D., & Joe, H. (2009). Generating random correlation matrices based on vines and extended onion method. *Journal of Multivariate Analysis*, *100*(9), 1989–2001. https://doi.org/10.1016/j.jmva.2009.04.008

Osnas, J. L. D., Katabuchi, M., Kitajima, K., Wright, S. J., Reich, P. B., Van Bael, S. A., Kraft, N. J. B., Samaniego, M. J., Pacala, S. W., & Lichstein, J. W. (2018). Divergent drivers of leaf trait variation within species, among species, and among functional groups. *Proceedings of the National Academy of Sciences of the United States of America*, *115*(21), 5480–5485. https://doi.org/10.1073/pnas.1803989115
